# Abstract sequential task control is facilitated by practice and embedded motor sequences

**DOI:** 10.1101/2020.02.06.937938

**Authors:** Juliana E. Trach, Theresa H. McKim, Theresa M. Desrochers

## Abstract

Everyday task sequences, such as cooking, contain overarching goals (completing the meal), sub-goals (prepare vegetables), and motor actions (chopping). Such tasks generally are considered hierarchical because superordinate levels (e.g., goals) affect performance at subordinate levels (e.g., sub-goals and motor actions). However, there is debate as to whether this hierarchy is “strict” with unidirectional, top-down influences, and it is unknown if and how practice affects performance at the superordinate levels. To investigate these questions, we manipulated practice with sequences at the goal and motor action levels using an abstract, or non-motor, task sequence paradigm (Desrochers et al., 2015; Schneider & Logan, 2006). In three experiments, participants performed memorized abstract task sequences composed of simple tasks (e.g., color/shape judgements), where some contained embedded motor response sequences. We found that practice facilitated performance and reduced control costs for abstract task sequences and subordinate tasks. The interrelation was different between the hierarchical levels, demonstrating a strict relationship between abstract task sequence goals and sub-goals and a non-strict relationship between sub-goal and motor response levels. Under some conditions, the motor response level influenced the abstract task sequence level in a non-strict manner. Further, manipulating the presence or absence of a motor sequence after learning indicated that these effects were not the result of an integrated representation produced by practice. These experiments provide evidence for a mixed hierarchical model of task sequences and insight into the distinct roles of practice and motor processing in efficiently executing task sequences in daily life.

## Introduction

Humans are remarkably adept at executing complex task sequences in daily life. Consider cooking dinner. Making dinner involves the execution of a number of subtasks (e.g., chop vegetables, boil water, etc.) in a particular order that contribute to the completion of an overarching task goal. Such goals can be considered hierarchical because they involve a superordinate goal (make dinner) that is subserved by task sub-goals (chop vegetables, boil water, add pasta), which, in turn, can be further broken down into motor actions (chopping, stirring, scooping). Thus, the completion of such hierarchical tasks involves processing at the goal, sub-goal, and motor levels, as well as coordinating across goal, sub-goal and motor levels.

Despite their complexity, humans execute hierarchical task sequences routinely in daily life. How does experience with such task sequences improve our ability to complete them? One clear possibility is that practice leads to improved performance of task sequences. Additionally, interactions between information at the goal, sub-goal, and motor levels could support efficient execution. Executing such hierarchical tasks may intuitively suggest a directionality of influence in that the overarching goal constrains the sub-goals, which in turn constrain the motor responses. However, it remains an open question, particularly in the context of abstract task sequences, how practice might improve performance and whether lower level responses might facilitate superordinate sub-goal and goal execution and representation.

We will use task sequences to address these questions and operationalize the kinds of hierarchical sequences we readily perform in daily life. Task sequences are events that contain an overarching *sequence-level* goal that dictates a series of individual *task-level* goals that each involve a *motor-level* response (goal, sub-goals, and motor sequences respectively in the previous example; Schneider & Logan, 2006; **Figure 1**). Additionally, we use the word “abstract” to emphasize that these task sequences are composed of a series of goals that require variable responses, rather than series of specific motor actions. This definition contrasts with previous work that uses “tasks sequences” to describe a series of discrete tasks with motor responses that occur in a random order over the course of a block of trials (e.g., Korb et al., 2017; Strobach et al., 2012) or series of motor actions. Task sequences in our framework are unique because they incorporate the aforementioned three levels into the hierarchy (**Figure 1**).

**Figure 1.**
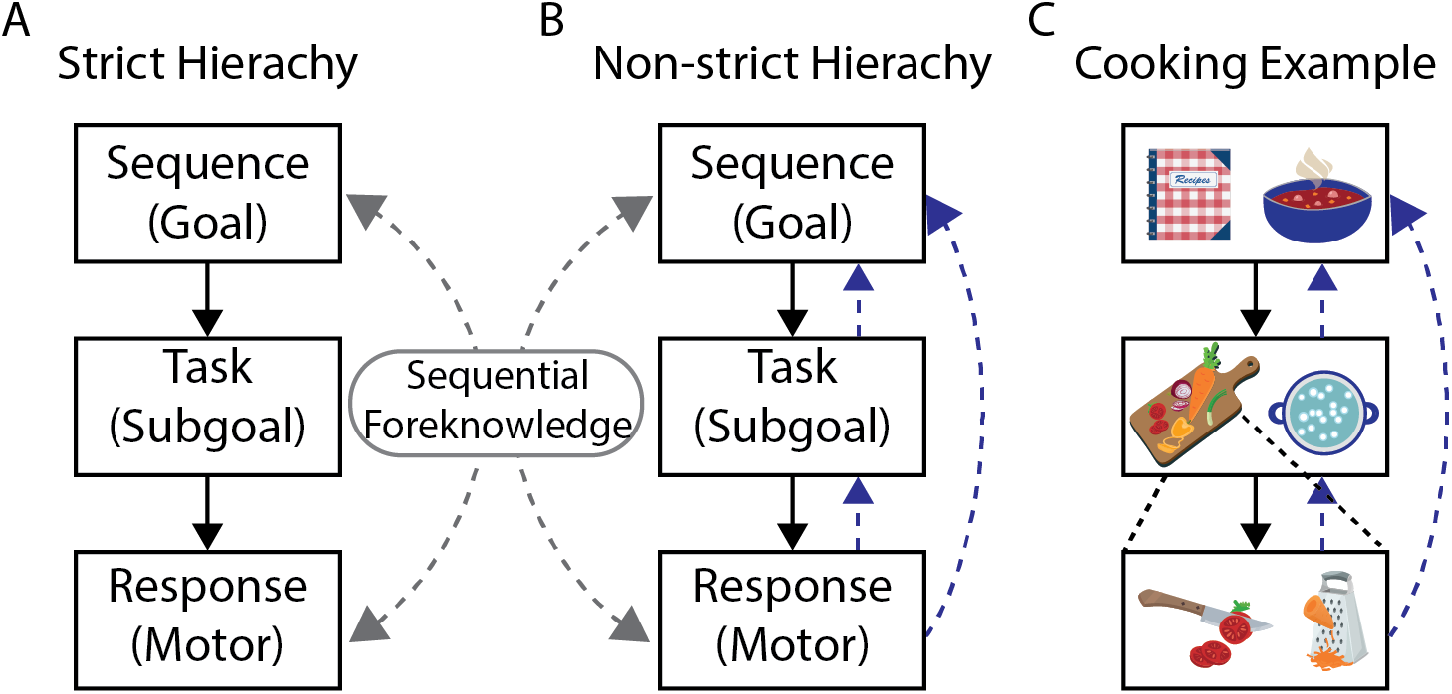
Example Hierarchy Schematic. (**A**) Example of strict hierarchy demonstrating unidirectional influence across levels. (**B**) Example of a non-strict hierarchy depicting multidirectional relationships between levels. (**C**) Cooking as an example of an abstract task sequence in daily life. Response (motor) level box shows examples of chopping and grating actions from the task (sub-goal level). Dashed gray lines indicate sequential foreknowledge. Dashed blue lines indicate predictions from experiments on the influence between levels. Image credit: Designed by macrovector/freepik.

Previous work has identified at least two possibilities for how task levels influence each other (Mayr & Bryck, 2005). One model is strictly hierarchical: higher order levels influence action selection at lower levels but not vice versa (**Figure 1A**). A strict hierarchical model would also exhibit a degree of modularity such that information within a level is relevant only at that stage of processing and does not interact with lower level selection. For example, the superordinate goal of making dinner (sequence level) constrains the relevant sub-tasks (task level), which in turn constrain the necessary motor responses (motor level), but the motor responses do not influence how the sub-task goals are processed. Alternatively, a non-strict hierarchical model could allow different levels to interact such that lower levels (e.g., motor responses) also influence higher-order levels (i.e. sequence and/or task level) during execution (**Figure 1B**) (Cock & Meier, 2013; Kikumoto & Mayr, 2020; Korb et al., 2017; Mayr & Bryck, 2005). In sum, a strict hierarchical model involves unidirectional flow of information such that superordinate levels influence subordinate levels, whereas a non-strict hierarchical model allows for bidirectional influences between multiple levels. Examining the directionality of information flow between sequence levels is essential to understanding how we execute such sequences efficiently in daily life.

In a series of three studies, we manipulated the information participants have about upcoming events, or *sequential foreknowledge,* to elucidate information flow between task levels. Specifically, we will examine the effect of increased foreknowledge at the sequence and motor levels on two behavioral indicators of sequential and hierarchical control described below. Within the context of task sequences, hierarchical control arises both from the relationship between sequence-level (goal) and task-level (sub-goal) information and from the relationship between task-level and motor responses in such tasks. We will use practice to increase sequential foreknowledge at the sequence level and examine practice effects on behavioral measures of hierarchical control. At the motor level, we will embed motor sequences into the button press responses that participants make to incorporate sequential foreknowledge. The strict and non-strict hierarchical models have distinct predictions for these manipulations. Broadly, the non-strict hierarchical model allows for manipulations at lower levels to affect processing at superordinate levels, while the strict model does not.

The two hierarchical models we have discussed have differentiable predictions about how the task and motor levels may, or may not, interact. Previous studies across at least two types of paradigms, task switching and dual-task, broadly support interaction between task and motor levels and therefore a non-strict hierarchy. Task switching paradigms measure switch costs, defined as increases in reaction times and error rates on trials where participants switch tasks (e.g. color to shape judgement) relative to those where they repeat tasks (for a review: Monsell, 2003). This behavioral measure can be used to examine the relationship between hierarchical levels, (B to C; **Figure 1**) and these costs are hypothesized to be due to the task-set reconfiguration necessary to switch tasks (Berryhill & Hughes, 2009; Draheim et al., 2016; Hirsch et al., 2018; Sabah et al., 2019; Strobach et al., 2018). A non-strict hierarchical model predicts that task switch costs would interact with switch costs at the motor level (i.e., the cost of switching over repeating a motor response) such that motor level switching would influence the task level, whereas a strict hierarchical model would predict no interactions. Studies have found interactions between task and response switching (Kikumoto & Mayr, 2020; Korb et al., 2017; Mayr & Bryck, 2005), supporting a non-strict hierarchy. These interactions have not been examined in the context of sequences of tasks.

In addition to task switching work, dual-task studies support the interaction between task and motor levels and a non-strict hierarchical interpretation. These studies focused on multiple streams of practiced sequences and showed that alignment of sequential content across levels affects learning (Cock & Meier, 2013; Deroost et al., 2007; Rah et al., 2000; Röttger et al., 2019; Weiermann et al., 2010; Weiermann & Meier, 2012; Zhao et al., 2019). These findings support a non-strict hierarchy between the motor and task levels in the context of practice and sequential information. However, these studies did not test for interactions indicative of non-strict hierarchical representation in the context of more abstract, task sequences. Thus, the following experiments were designed to test the interaction of task and motor levels and the effects of practice within the context of hierarchical control necessary for sequential behavior.

Hierarchical direction of influence can also be examined at a more superordinate level, i.e., the abstract sequence or goal level. Analogous to switch costs, initiation costs are increases in reaction times at the beginning of abstract sequences, relative to subsequent positions in the sequence (Desrochers et al., 2015; Farooqui & Manly, 2019; Schneider & Logan, 2006). These initiation costs provide evidence for hierarchical control between the sequence and task levels (A to B, **Figure 1**; Desrochers et al., 2015; Farooqui, Mitchell, Thompson, & Duncan, 2012; Sternberg et al., 1988). The presence of initiation costs supports a strict hierarchy; however, the presence or absence of non-strict influences have not been explicitly tested at the abstract task sequence level, with or without practice. Broadly, because practice reduces, but does eliminate switch costs which are thought to reflect similar processes at the task level, (Berryhill & Hughes, 2009; Stoet & Snyder, 2007; Strobach et al., 2012), we predict that practice will reduce initiation costs. We will also probe if and how lower levels influence and interact with superordinate levels and how practice affects these potential relationships by examining the effects on initiation costs.

The goal of our studies is to investigate the independence or interactions between the task and motor response levels in the context of abstract task sequences. We operationalized the sequence, task and motor levels of a behavioral task based on a switching tasks in sequences paradigm developed to study hierarchical control (Desrochers et al., 2015; Schneider & Logan, 2006). We manipulated sequential foreknowledge, at the sequence level via practice and at the motor level via embedded motor sequences and assessed the effect of this manipulation on behavioral manifestations of control at the task level in switch costs, and at the sequence level in initiation costs. Crucially, if embedded motor sequences reduce control costs at superordinate levels, this result would provide evidence that lower levels in the hierarchical task influence superordinate levels in a non-strict, rather than strict, hierarchical manner. Overall, we found practice reduced behavioral costs associated with sequence-level processing but not task-level processing. Additionally, we found that motor-level foreknowledge reduced these costs at the task level but evidence for embedded motor sequences affecting sequence level processing was less consistent. Our findings point to a mixed hierarchical model where some levels exhibit a strict hierarchical organization and others exhibit non-strict relationships in the execution of task sequences.

## Experiment 1

The goal of Experiment 1 was to examine the effects of increased sequential foreknowledge at the superordinate sequence level of a hierarchical task sequence paradigm. We modified a task previously used to examine the behavioral and neural correlates of hierarchical cognitive control in the execution of abstract task sequences (Desrochers et al., 2015; Schneider & Logan, 2006). In this task, participants executed abstract task sequences, each comprised of five individual tasks: simple color or shape judgements. We operationalized an increase of sequential foreknowledge at the superordinate sequence level by having participants practice two specific abstract task sequences. In the test phase participants executed the practiced (Familiar) sequences and new abstract task sequences that they had not seen during the practice phase (Novel). Importantly, the constituent simple task judgements were the same between Familiar and Novel sequences, but the order was either practiced or new. We then compared performance on Familiar and Novel sequences to examine how sequential foreknowledge affected performance.

Overall, we hypothesized that increased sequential foreknowledge would facilitate performance on abstract task sequences. In particular, we aimed to adjudicate between a strict and non-strict hierarchical model of task sequence execution by determining if increased foreknowledge had specific effects on initiation or switch costs. A specific effect on initiation costs would suggest that sequential foreknowledge at the sequence level reduces the control costs at the top level of this hierarchical task associated with initiating and executing a task sequence, supporting a strict hierarchical model. In contrast, overall reductions in RT would support a non-strict hierarchical model. At the task level, a reduction in switch costs due to practice would suggest that increased foreknowledge at the sequence level yields task level improvements as well, providing evidence for a non-strict hierarchical model. Alternatively, if there is no reduction in switch costs with increased practice, that would suggest independence between levels and provide evidence for a strict, modular hierarchical model.

## Methods

### Participants

Twenty-nine (*n* = 21 female) adults between the ages of 18-35 (*M* = 20 *SD* = 1.7) participated in the study. All participants were right-handed, had normal or corrected-to-normal vision, and reported that they were not colorblind. Individuals with neurological or psychiatric conditions, brain injury, or reported use of psychoactive medications or substances were excluded from participating. Participants were recruited from the Brown University campus and the surrounding community as well as from the student course credit participant pool (administered through Sona systems). Participants were compensated for their time ($10/hour) or received course credit for the approximately 1-hour study. All participants gave informed, written consent as approved by the Human Research Protections Office at Brown University.

### Procedure

Trials: On each trial of the abstract sequential task, participants were presented with a stimulus and were asked to respond to one of three features of the presented image: size, color, or shape. The background screen was grey, and all text was presented in white font. After the presentation of the stimulus, the participant had four seconds to respond with a button press based on the relevant stimulus feature (Desrochers et al., 2015). The stimulus remained on the screen until the response was made or the four seconds elapsed without response, giving participants ample time to respond. After the response period, a black fixation cross appeared centrally during an intertrial interval (ITI) of 250 ms. All ITIs were of this same duration. No feedback was given after each button press.

Each stimulus feature had multiple options. Size was large (7.0 × 7.0 cm) or small (3.5 × 3.5 cm); shape was circle, square, triangle, or star; and color was red, green, pink, or blue. During the practice phase, only two of the four stimulus feature options were used for the color and shape choices (e.g., blue/red circles/triangles), then the remaining two (e.g., green/pink squares/stars) were used during the testing phase. This procedure ensured that effects of learning evident in the experimental phase could not be attributed to familiarity with the specific stimuli, but rather to familiarity with the sequence of feature judgements. Shape and color combinations were counterbalanced across participants.

Participants used the ‘j’ and ‘k’ keys to select a response with the index and middle finger of their right hand. Feature-response mappings changed between the practice and test phases of the experiment (along with the stimuli) and were counterbalanced across participants. Reminders of all three feature-response mappings were presented with each stimulus at the bottom of the screen for all trials (**Figure 2A)**.

**Figure 2.**
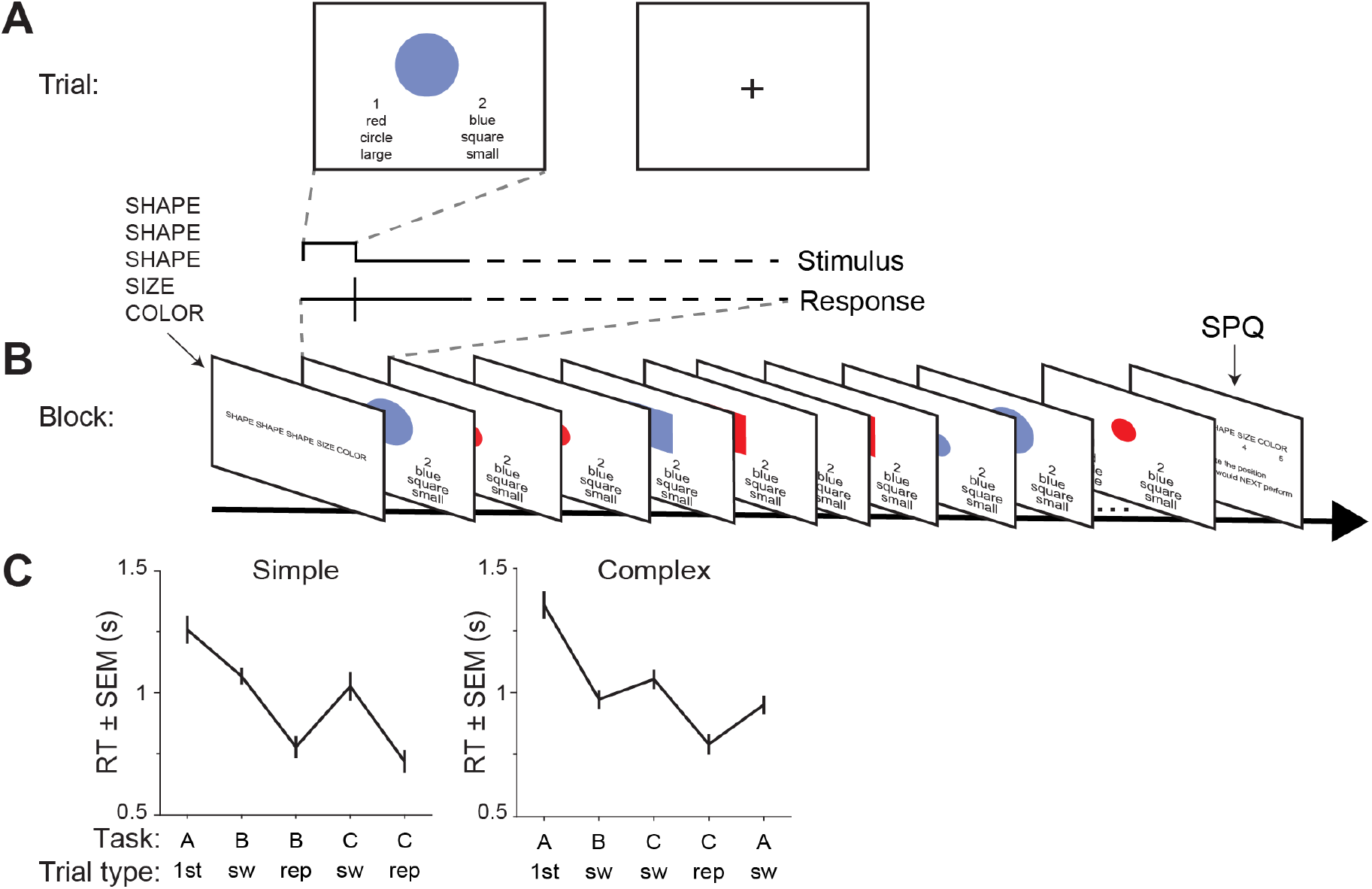
Experiment 1 Task Paradigm. (**A**) Example of task trial. Participants were instructed to remember a five-item sequence at the beginning of the block (4s) and make the appropriate feature judgement on each trial. The stimulus remained on the screen until a response was made (max 4 s). (**B**) Trials were structured into blocks that ended with a sequence position question (SPQ) to probe the participant to respond what the next image in the sequence would have been if the trial continued. (**C**) Example reaction time (RT) profiles for a simple and complex sequence. The three stimulus features (color, shape, and size) have been generalized as the letters “A”, “B”, and “C” such that instances of the same structure within different specific task sequences (e.g., color, shape, size, size, size; and size, color, shape, shape, shape) are averaged together. 1^st^, first; sw, switch; rep, repeat.

Blocks: Participants knew which stimulus features were relevant on each trial due to the instruction presented at the beginning of each block of trials. At the start of each block, a 5-item feature sequence was displayed (5 seconds; s), e.g., “shape, shape, shape, size, color”. Participants were instructed to remember the sequence and respond to the stimuli as they appeared on each trial by following the order of this instructed sequence. For example, in the sequence “shape, shape, shape, size, color”, participants responded to the stimulus feature of shape of the first, second, and third stimulus, the size of the fourth, and the color of the fifth stimulus (**Figure 2B**). Participants repeated the abstract task sequence until the end of the block, which contained 15-19 trials. There were no external cues provided, other than at the beginning of the block, to indicate the position within the abstract task sequences. Therefore, the position in the sequence was tracked internally by participants. The order of the stimuli within the sequence, and consequently, the correct key press responses for each sequence, were randomized such that there were no predictable motor sequences embedded in the responses.

The block of trials could terminate at any position within the sequence with equal frequency. At the end of each block, participants answered a sequence position question. Participants responded using five keys (‘J’, ‘K’, ‘L’, semicolon, or apostrophe) to indicate what the next feature judgment in the sequence would have been, had the sequence continued. This question was used to encourage participants to continue to execute the abstract task sequence as instructed throughout the block and not chunk or rearrange the elements such that they were grouped differently than originally instructed. The question also served as an indicator of whether they may have lost their place in the sequence during the block. Participants had five seconds to make a response to the sequence position question at the end of the block. Once participants responded, a fixation cross appeared (2 s) before the start of the next block.

Sequence complexities: Sequences were classified as either simple or complex (**Table 1**). Simple sequences contained two “internal” (positions 2, 3, 4, or 5) task switches (e.g., “shape, <u>color</u>, color, <u>size</u>, size”; task switch trials underlined), and complex sequences contained three internal task switches (e.g., “shape, color, size, size, shape”). Although these sequence types differed in the number of internal task switches, the total number of switches and repeats were balanced across sequence types when sequences were repeated throughout the block. In simple sequences, the first feature judgment in the sequence was a task switch when restarting the sequence (i.e. position 5 to position 1), and in complex sequences the first feature judgment in the sequence was always a repeat trial (**Figure 2C**). The primary purpose of the different sequence complexities was to ensure that abstract task sequences began with both switch and repeat trials. This counterbalancing was only possible by varying the number of internal task switches, given the constraints we used to construct task sequences (e.g., an equal number of switches and repeats for each sequence).

**Table 1.** Sequence structures and example sequences.

| Structure | Example Sequence of Tasks |
| --- | --- |
| Simple |  |
| AAABC | Color, Color, Color, Size, Shape |
| AABCC | Size, Size, Shape, Color, Color |
| AABBC | Shape, Shape, Size, Size, Color |
| ABBBC | Size, Color, Color, Color, Shape |
| ABBCC | Shape, Color, Color, Size, Size |
| ABCCC | Color, Shape, Size, Size, Size |
| Complex |  |
| AABCA | Shape, Shape, Size, Color, Shape |
| ABBCA | Color, Size, Size, Shape, Color |
| ABCCA | Size, Color, Shape, Shape, Size |
| ABC AA | Shape, Size, Color, Shape, Shape |

Runs: In a session, participants first completed two short training blocks to introduce them to the feature judgments and stimuli (10-14 trials). Then, the practice phase consisted of four runs of six blocks each where participants performed two abstract task sequences, one simple and one complex. These two practiced sequences are referred to as Familiar sequences. For the test phase, participants executed another four runs of six blocks each that contained sequences that they had not seen before (Novel) intermixed with the Familiar sequences practiced previously (**Figure 2C**). Novel sequences were selected such that they did not have the same pattern of switch and repeat trials as practiced sequences. For example, a Familiar sequence that was “color, size, shape, shape, shape” follows a general “A, B, C, C, C” task structure, and so the sequence “shape, color, size, size, size” would not be used as a comparable Novel sequence **(Table 1)**. This design avoided any transfer of learning effects from familiar sequences that might occur due to sequence structure. Participants initiated each run by pressing the spacebar. Upon task completion, participants filled out an online (Qualtrics), post-test questionnaire with questions about their experience with the task. The full experiment lasted approximately one hour. The task was programmed in Psychophysics Toolbox (Brainard, 1997; http://psychtoolbox.org/; RRID:SCR_002881) and run in Matlab (MathWorks; RRID:SCR_001622).

### Analysis

All analyses across all experiments were conducted in Matlab (MathWorks; RRID:SCR_ 001622). No participants had an error rate greater than our exclusion criterion of 20%, as in previous studies (Desrochers et al., 2015). The first iteration of the sequence (five trials) in each block were excluded from analyses to avoid any confounding block initiation effects. We also excluded trials with a reaction time of less than 100 ms, as a reaction time of less than 100 ms is not sufficient to perform the task as instructed by first perceiving the stimulus and then making the appropriate judgement. Trials with a reaction time of 4 seconds were also excluded as that indicated that the trial had elapsed without the participant making a response.

Reaction times (RTs) and error rates (ERs) were submitted to repeated measures ANOVAs (rmANOVAs) and t-tests where appropriate. Simple and complex sequences were combined for statistical testing, as the focus of our experiments was on features that were common to both sequence types. Simple and complex sequences contain differing numbers of switch and repeat trials at noninitial positions. Therefore, to avoid weighting the mean of noninitial positions, we calculated an unweighted mean for noninitial positions using the following procedure. For each participant, all switch trials in noninitial positions and all repeat trials in noninitial positions were first separately averaged. Then an unweighted mean of these two averages was computed to find the mean of the noninitial positions. Our primary, planned analyses were rmANOVAs to evaluate sequence initiation across conditions and therefore contained factors for condition (Familiar and Novel) and trial type (first and noninitial). Additionally, we planned to evaluate practice effects on switch costs with a rmANOVA with factors for condition (Familiar and Novel) and trial type (switch and repeat). We further tested whether practice (Familiar, Novel) affected trial type (first, noninitial) based upon switch type (switch and repeat). When sphericity assumptions were violated, we used Greenhouse Geisser correction for the degrees of freedom.

## Results

Overall, participants readily learned the task sequences, and results replicated previous studies. Participants performed the five-item sequences well at test (error rate, ER: *M* = 2.9%, *SD* =2.6%) and were faster at Familiar sequences during the test phase as compared to the practice phase (practice RT: *M* = 1.1 s, *SD* = 0.22 s; test RT: *M* = 0.96 s, *SD* = 0.17 s; **Table 2, Figure 3A**). ERs did not differ between practice and test (*t*_28_ = 1.1, *p* = 0.28, *d* = 0.27, **Figure 3B**). Different sequence complexities were included solely to counterbalance task switching and repeating at each sequence position. While complex sequences were slower than simple sequences (*F*_0.58,16_ = 11, *p* = 0.0022, η_p_^2^= 0.29), there were no interactions between complexity and trial type (*F*_1.2,32_ = 0.64, *p* = 0.53, η_p_^2^= 0.022) or condition (*F*_0.58,16_ = 0.0064, *p* = 0.94, η_p_^2^= 0.00023). Therefore, for all following analyses, we combined across sequence complexities. Initiation costs were calculated by creating an unweighted mean of noninitial positions for comparison with the first serial position in sequences (see Methods). The presence of initiation costs in RT and not ER (**Table 2**) confirmed that participants were performing the judgements as sequences and replicated previous results (Desrochers et al., 2015, 2019; Schneider & Logan, 2006). Because this key sequential indicator is present in RT, subsequent analyses focus on RT measures to determine the effects of practice on the hierarchical control costs.

**Figure 3.**
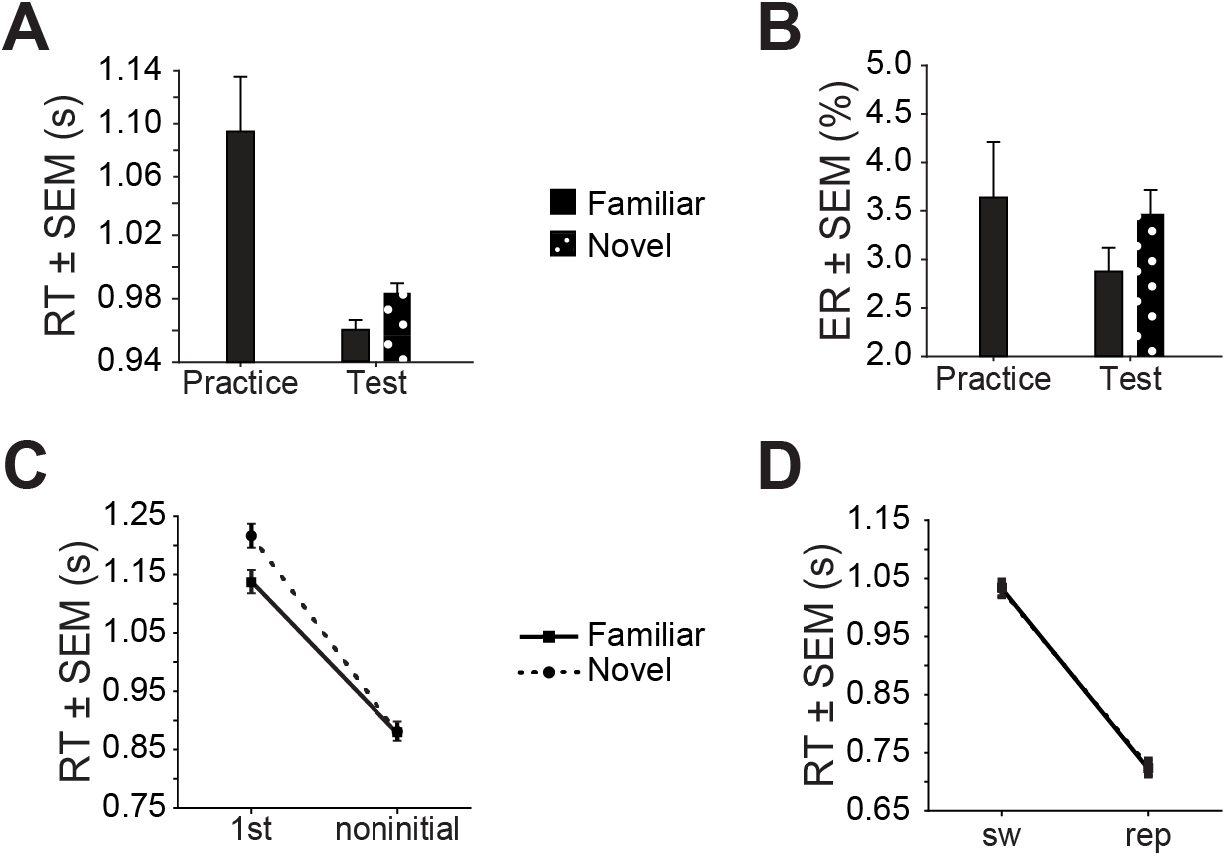
Experiment 1 Results. (**A**) Average reaction time (RT) plotted for practice and test study phases by condition (Familiar vs. Novel). (**B**) Average error rate (ER) as in A. (**C**) Plot of RT for first (1st) versus noninitial (unweighted mean for noninitial positions) trial types by condition. (**D**) Average RT plotted for switch (sw) and repeat (rep) trials by condition. F, Familiar/solid/squares; N, Novel/dotted/circles.

**Table 2.** Experiment 1 t-tests comparing 1st and noninitial positions for RT and ER.

| Condition | Reaction Time |  |  |  | Error Rate |  |  |
| --- | --- | --- | --- | --- | --- | --- | --- |
|  | *dfs* | *t* | *p* | *d* | *t* | *p* | *d* |
| Familiar | 28 | 9.1 | <0.001 | 1.36 | 0.46 | 0.64 | 0.074 |
| Novel | 28 | 13 | <0.001 | 1.74 | -0.85 | 0.41 | 0.21 |

To examine the main question of how practice at the sequence level affects performance at the sequence and task levels, we compared Familiar and Novel sequences. If sequence and task levels exhibit a strict hierarchical relationship, we expect practice with sequence-level information to specifically reduce initiation costs. In contrast, a non-strict hierarchical model allows for practice with sequence-level information to affect task level processing as well and would not predict a specific reduction in initiation costs. Importantly, in this comparison we isolated effects at the task sequence level as all other levels have been practiced equally (i.e., individual task judgements and motor responses). We found that practice decreased initiation costs in Familiar sequences (**Figure 3C**, **Table 3**, **Table 4**; interaction: *F*_0.74,21_ = 5.2, *p* = 0.03, η_p_^2^= 0.16), specifically at the first position in the sequence (post-hoc t-test, Bonferroni-adjusted a = 0.025: first position, *t*_28_ =-2.4, *p =* 0.024, *d* = 0.35; noninitial positions, t_28_ =-0.060, *p =* 0.95, *d* = 0.0042). These results suggest that abstract task sequence practice selectively affects sequence-level performance in a strict hierarchical manner.

**Table 3.** Experiment 1 rmANOVA for RT and ER initiation cost.

| Factor | Reaction Time |  |  |  | Error Rate |  |  |  |
| --- | --- | --- | --- | --- | --- | --- | --- | --- |
| | *dfs* | *F* | *p* | $\eta_p^2$ | *dfs* | *F* | *p* | $\eta_p^2$ |
| Condition | 0.74, | 5.0 | 0.03 | 0.15 | 0.79, | 0.17 | 0.68 | 0.0061 |
| (Familiar, Novel) | 21 |  |  |  | 22 |  |  |  |
| Trial type<br>(1<sup>st</sup>, noninitial) | 0.74,<br>21 | 185 | <0.001 | 0.87 | 0.79,<br>22 | 0.21 | 0.65 | 0.0075 |
| Condition ×<br>Trial type | 0.74,<br>21 | 5.2 | 0.03 | 0.16 | 0.79,<br>22 | 0.97 | 0.33 | 0.033 |

**Table 4.** Experiment 1 rmANOVA for RT and ER initiation cost.

| Condition | Trial type |  |
| --- | --- | --- |
|  | First | Noninitial |
| Familiar |  |  |
| RT (s) | 1.1 (0.041) | 0.88 (0.030) |
| ER (%) | 2.9 (0.63) | 2.7 (0.49) |
| Novel |  |  |
| RT (s) | 1.2 (0.041) | 0.88 (0.030) |
| ER (%) | 2.7 (0.64) | 3.4 (0.46) |

Standard error of the mean (SEM) appears in parentheses.

Further support for a strict hierarchical interpretation is provided by examining the influence of practice on other control costs: switch costs. A strict hierarchical account would not predict reductions in switch costs due to sequence-level practice whereas a non-strict hierarchical model would. Switch costs in noninitial positions were not reduced in Familiar sequences (interaction: *F*_0.62,21_ = 0.0036, *p* = 0.95,η_p_^2^= 0.00013, **Figure 3D, Table 5**). Further, practice affected first position trials equivalently, regardless of whether the first position trial was a task switch or task repeat (condition × trial type × switch type: *F*_0.64,0.64,18_ = 1.2, *p* = 0.28, η_p_^*2*^= 0.042; **Table 6**). These results suggest that practice selectively facilitated the execution of abstract task sequences at sequence initiation without reducing control costs at the task level, thus supporting a strict hierarchical model.

**Table 5.** Experiment 1 rmANOVA for RT switch cost.

| Factor | Reaction Time |  |  |  |
| --- | --- | --- | --- | --- |
| | *dfs* | F | *p* | $\eta_p^2$ |
| Condition<br>(Familiar, Novel) | 0.62,<br>21 | 0.099 | 0.76 | 0.0035 |
| Trial type<br>(sw, rep) | 0.62,<br>21 | 196 | <0.001 | 0.87 |
| Condition ×<br>Trial type | 0.62,<br>21 | 0.0036 | 0.95 | 0.00013 |

**Table 6.** Experiment 1 three-way rmANOVA with condition, trial type and switch type factors for RT.

| Factor | Reaction Time |  |  |  |
| --- | --- | --- | --- | --- |
| | *dfs* | F | *p* | $\eta_p^2$ |
| Condition<br>(Familiar, Novel) | 0.64,<br>18 | 4.8 | 0.037 | 0.15 |
| Trial type<br>(1<sup>st</sup>, noninitial) | 0.64,<br>18 | 187 | <0.001 | 0.87 |
| Switch type<br>(sw, rep) | 0.64,<br>18 | 38 | <0.001 | 0.57 |
| Condition ×<br>Trial type | 0.64,<br>18 | 5.1 | 0.033 | 0.15 |
| Condition ×<br>Switch type | 0.64,<br>18 | 0.67 | 0.42 | 0.023 |
| Trial type × | 0.64, | 74 | <0.001 | 0.72 |
| Switch type | 18 |  |  |  |
| Condition × | 0.64, | 1.2 | 0.28 | 0.042 |
| Trial type × | 0.64, |  |  |  |
| Switch type | 18 |  |  |  |

Results from this experiment support a strict hierarchical relationship between sequence and task level information, but open questions remain. First, it is possible that additional practice could enact processes that improve performance across all levels of the abstract task sequences in a non-strict manner. Second, rarely in daily living are abstract task sequences isolated from the motor sequences that subserve them. Therefore, interactions between the motor and superordinate task and sequence levels (**Figure 1B**) also need to be examined for their strict or non-strict hierarchical properties. Both of these questions will be addressed in Experiment 2.

## Experiment 2

The aim of Experiment 2 was to determine if increased sequence-level practice and motor-level sequential foreknowledge changed the relationship between levels of a hierarchical task sequence. To accomplish this goal, we modified the same abstract sequence task used in Experiment 1. First, we expanded practice at the sequence level by approximately tripling the number of practice trials and spreading practice between two experimental sessions on separate days. Second, we introduced the Motor Familiar condition that incorporated sequential foreknowledge at the motor level. This novel manipulation at the lowest level of the hierarchy (**Figure 1B**) in the context of a more abstract, task sequence paradigm further tested hypotheses regarding a strict versus non-strict relationship across hierarchical levels in the context of sequences that more closely approximate the complex structure of daily life.

## Methods

### Participants

Recruitment, inclusion criteria, and consenting procedures were the same as in Experiment 1. Thirty-five (*n* = 21 female) individuals participated in the study. Individuals were excluded for neurological or psychiatric conditions, brain injury or using psychotropic medications or substances (*n* =3); ER greater than 20% (*n* = 1); and not completing both sessions of the experiment (*n* = 3). Thus, twenty-eight (*n* = 16 female) adults between the ages of 18-35 (*M* = 22, *SD* = 4.2) were included in analyses.

### Procedure

We adapted the task used in Experiment 1 to examine the central hypotheses of Experiment 2. The general structure of trials and blocks was the same as in Experiment 1, with the following modifications. We increased the number of possible responses on each trial from two to three (‘J,’ ‘K’ or ‘L’ keys). We did this in order to increase the number of possible embedded motor sequences (see below). We also replaced the size feature judgment with a texture feature judgment (solid, dotted, or striped) to create three easily distinguishable options for each task feature (color: red, green, or blue; shape: square, circle, or triangle; **Figure 4A-B**). Finally, we used a two-session format in Experiment 2 to accommodate new condition types and to increase the number of trials participants completed with Familiar sequences.

**Figure 4.**
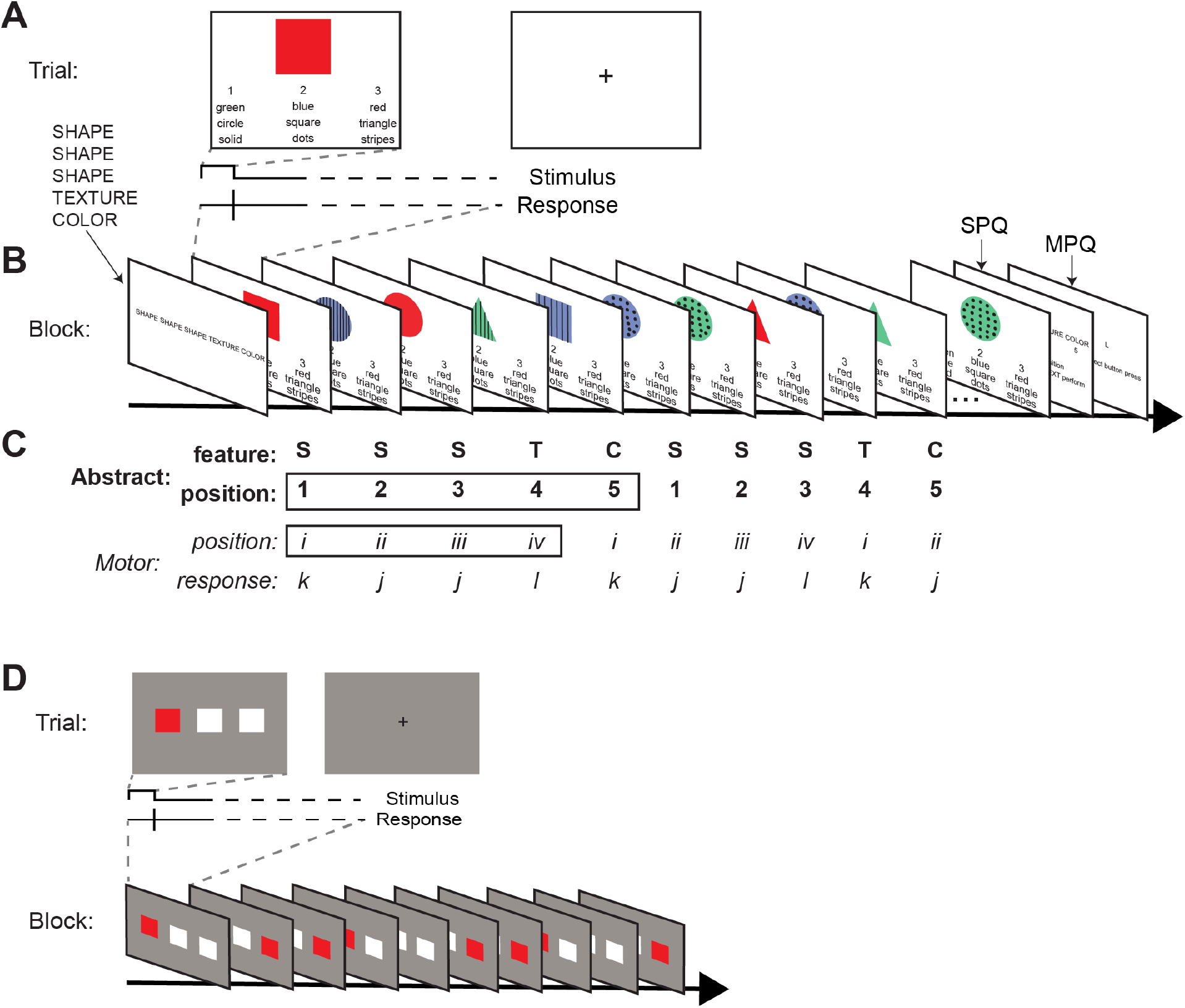
Experiments 2 & 3 Task Paradigm. (**A**) Task design used in Experiments 2 and 3 that contains abstract and motor sequences. (**B**) The trial and block structure were similar to the structure in Experiment 1, except that there were three possible button press responses (instead of two). (**C**) Two example iterations of a simple sequence with an embedded motor sequence. Each trial is classified by stimulus feature (S, shape; T, texture; C, color), abstract position (1-5), motor position (i-iv) and motor response (j,k,l). Note that the first position in the abstract sequence is also a switch trial (change from C to S judgement). Motor sequences have four items and abstract sequences have five items therefore resulting in a “frame shift” where there is no consistent relationship between the abstract and motor sequences. (**D**) Example Serial Reaction Time Task (SRTT) trial and block. Participants were instructed to press the key that spatially corresponds (‘J’, ‘K’, or ‘L’ key) to the location of the red square on each trial.

Motor sequences: Four-member motor sequences of button press responses were embedded in one condition, Motor Familiar. Participants were not given explicit instruction about the presence or identities of these sequences; therefore, any knowledge of the motor component was gained implicitly. An example embedded motor sequence was ‘KJJL’. All motor sequences had one repeat trial (position 3 in this example). The presence of a repeat trial in the motor sequence means that one button is pressed with greater frequency in Motor Familiar blocks. We addressed this fact by including two different motor sequences for each participant, and with additional analyses presented in the results section.

The frame shift between the four-item motor sequences (e.g. i ii iii iv) and the five-item abstract task sequences (e.g. 1 2 3 4 5) avoided creating a predictable sequence of features (e.g. 1/i, 2/ii, 3/iii, 4/iv, 5/i – 1/ii, 2/iii, 3/iv, 4/i, 5/ii, etc.; **Figure 4C**). Without this frame shift, for example, if the first position in the example abstract task sequence (e.g. “shape”) was always paired with the first position in the motor sequence (‘K’), the stimulus that would appear on first position trials would always be a square, and the same would be true of all positions in the abstract task sequences. Due to the offset of the abstract and motor sequences, a participant had to complete four iterations of the abstract sequence in order to execute all possible combinations of abstract and motor sequence position trials, making unlikely that they memorized the series of responses. To further reduce the likelihood of memorization, participants did not perform the same abstract sequence in two consecutive blocks.

All blocks (including conditions Familiar, Motor Familiar, and Novel) were 45-49 trials in order to accommodate the new Motor Familiar condition and ensure a sufficient number of each possible alignment of positions in the motor and abstract sequences. Each participant practiced two Motor Familiar sequences and two Familiar sequences during session 1. Those abstract task sequences that did not contain an underlying motor sequence had the key press responses randomized such that there was no repeating pattern.

Blocks: At the end of each block, in addition to the Sequence Position Question, we introduced a button guess judgment. Participants were asked to guess what the next correct button press response (‘J’, ‘K’, or ‘L’) would have been had the block continued. The button guess judgment was used to assess awareness of the motor sequences. In Motor Familiar sequences, participants might have been able to predict the next correct button press if they gained some knowledge of the embedded motor sequences. This question was included at the end of each block, regardless of whether there was an embedded motor sequence. Participants had five seconds to respond to each of these questions.

Sessions: Participants completed two sessions on separate days. In the first session, participants were first familiarized with each feature judgment individually. Participants then executed two training blocks of 10-15 trials (2-3 abstract task sequences) each that were not included in analyses. Then, they completed six total runs (8 blocks each) of practice. Each block contained one of the possible combinations of complexity and conditions: a simple or complex Familiar, or a simple or complex Motor Familiar. The order of the blocks was counterbalanced across runs and participants.

The second session occurred within two days (*M* = 1.8, *SD* = 0.87) of the first session. During the test session, participants executed the sequences they had practiced during session one (Familiar and Motor Familiar), with the addition of Novel simple and complex sequences. The second session also contained six runs of eight blocks each. There were 16 blocks of each of the Motor Familiar sequences and eight blocks of each of Familiar and Novel sequences. We included more blocks of Motor Familiar sequences to obtain sufficient trials of the combination of abstract and motor sequence positions. Each session lasted approximately 1.5 hours.

At the end of each session, participants completed an online post-test questionnaire with questions about their experience with the task. We also used the post-test questionnaire to assess awareness of the embedded motor sequences. Importantly, questions at the end of session one obliquely alluded to the presence of a motor sequence by asking if any sequences “seemed easier” than others, as well as whether participants had intuitions about such occurrences. After the second session, participants were directly asked if they had noted any consistencies in the series of button press responses they executed. Example questions were: “Did you notice any pattern to the sequence of buttons you pressed to respond to the sequence?” (yes, maybe, no), and “If so, how sure are you that you noticed a pattern?” (1, very unsure to 4, certain). Participants were also asked to reproduce any button press consistencies they identified.

Serial reaction time task: A SRTT (Nissen & Bullemer, 1987) was included at the end of session two (after the post-task questionnaire) to examine the execution of the motor sequence without the abstract component (**Figure 4D**). On each trial, three squares appeared on the screen. Participants had to press the key (‘J,’ ‘K,’ or ‘L’) that corresponded to the position of the red square (the other two were white, **Figure 4D**). Participants initiated a block of trials by pressing the spacebar, and there were no time limits on trials. The red square remained red until the participant correctly responded to its spatial location, and then that square turned white and the next square immediately turned red. Participants completed 6 blocks (25 trials each) of the SRTT. The order of the blocks was counterbalanced across participants. In two blocks, the order of the red square’s location was random. In the other four blocks, the sequence of red square locations was the same as the four-item sequence of button presses embedded in the Motor Familiar condition from the abstract sequence task.

### Analysis

RTs and ERs were submitted to rmANOVAs and t-tests where appropriate. To test for effects of practice, we again conducted condition (Familiar, Novel) × trial type (first, noninitial or switch, repeat) rmANOVA comparisons between Familiar and Novel conditions. Across Experiments 1 and 2, we tested for differences in the effect of practice by including a group factor (Experiment 1 or 2) in the condition × trial type rmANOVA. To further test the effects of adding an embedded motor sequence, we tested planned comparisons between Motor Familiar and Novel conditions as well as Motor Familiar and Familiar conditions. We conducted control analyses addressing motor response frequency (repeat or switch) between conditions (Motor Familiar, Familiar). We additionally tested for congruency effects between the motor level (switch or repeat) and the task level (switch or repeat) in the Motor Familiar condition. Further, we tested whether practice affected trial type (first, noninitial) based upon switch type (switch and repeat) as in Experiment 1. As in Experiment 1, our primary comparisons either compare RTs on repeat trials to RTs on switch trials or RTs on first position trials to the unweighted average RT of noninitial positions (see Experiment 1 methods). As planned, we combined across simple and complex sequences after verifying there were no significant differences in performance. When sphericity assumptions were violated, we used Greenhouse Geisser correction for the degrees of freedom.

## Results

We replicated the basic results from Experiment 1 in Experiment 2. Participants again performed the task well (test ER: *M* = 6.9%, *SD* = 4.6%). General performance on the practiced conditions improved as illustrated by faster RTs at test than at practice (average Familiar RT at practice versus test: *t*_27_ = 8.5, *p* < 0.001, *d* = 1.2; average Motor Familiar RT at practice versus test: *t*_27_ = 8.5, *p* < 0.001, *d* = 1.3; **Figure 5A**). There were no accuracy differences between practice and test sessions in the practiced conditions (average Familiar ER at practice versus test: *t*_27_ = 0.63, *p* = 0.53, *d* = 0.14; average Motor Familiar ER at practice versus test: *t*_27_ = 1.08, *p* = 0.29, *d* = 0.27; **Figure 5B**). All subsequent analyses presented were from the test phase and combine simple and complex sequences as there were no significant differences between complexities (*F*_0.57,15_ = 0.23, *p* = 0.63, η_p_^2^= 0.0086). Initiation costs were observed across conditions (Familiar, Motor Familiar, and Novel) in RT and not in ER (**Table 7**), replicating previous experiments (Desrochers et al., 2015; Schneider & Logan, 2006) and Experiment 1 results.

**Figure 5.**
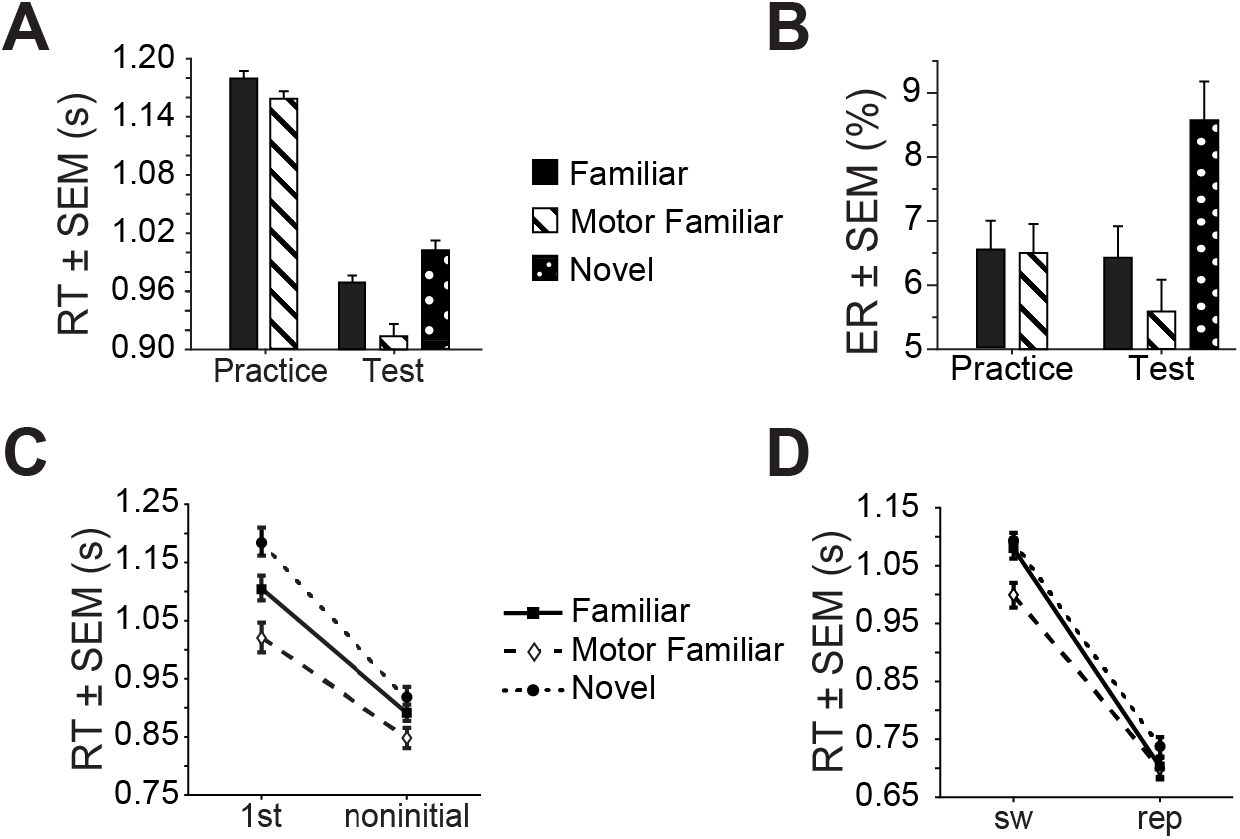
Experiment 2 Results. **(A)** Average reaction time (RT) plotted for practice and test study phases by condition (Familiar, Motor Familiar, and Novel). **(B)** Average error rate (ER) as in A. **(C)** Plot of RT for 1^st^ versus noninitial trial types by condition. **(D)** Plot of RT for switch (sw) and repeat (rep) trial types by condition. Familiar/solid/squares; Motor Familiar/dashed/diamonds; N, novel/dotted/circles.

**Table 7.** Experiment 2 t-tests comparing 1st and noninitial positions for RT and ER.

| Condition | *dfs* | Reaction Time |  |  | Error Rate |  |  |
| --- | --- | --- | --- | --- | --- | --- | --- |
|  |  | *t* | *p* | *d* | *t* | *p* | *d* |
| Familiar | 27 | 6.5 | <0.001 | 1.1 | 1.3 | 0.21 | 0.19 |
| Motor Familiar | 27 | 4.7 | <0.001 | 0.83 | 1.5 | 0.15 | 0.19 |
| Novel | 27 | 8.5 | <0.001 | 1.4 | -1.2 | 0.24 | 0.15 |

To determine if additional practice could enact processes that improve performance across all levels of the abstract task sequences in a non-strict manner, we first performed planned comparisons between Familiar and Novel sequences. As in Experiment 1, evidence that practice with sequence-level information specifically affected initiation costs would support a strict hierarchical model, whereas a general reduction in RT would support a non-strict hierarchical account. We replicated our results from Experiment 1 at the sequence level. RTs were faster for Familiar sequences (*F*_0.60,16_ = 9.3, *p* = 0.0051, η_p_^2^= 0.26), and practice differentially affected the first as compared to subsequent positions (*F*_0.60,16_ = 5.3, *p* = 0.030, η_p_^2^= 0.16; **Figure 5C**; **Table 8** and **Table 9**). The reduction in initiation cost may be driven by a decrease at the first position (post-hoc t-test, Bonferroni-adjusted a = 0.025: first position, *t*_27_ = 2.9, *p* = 0.0071, *d* = 0.34; noninitial positions, *t*_27_ = 2.1, *p* = 0.046, *d* = 0.17), and these effects were not significantly different from those in Experiment 1 (condition [Familiar, Novel] × trial type [first, noninitial] × experiment [Experiment 1, Experiment 2] rmANOVA: experiment: *F*_0.69,38_= 0.0077, *p* = 0.93; experiment × condition: *F*_0.69,38_ = 0.23, *p* = 0.63; experiment × trial type: *F*_0.69,38_ = 2.4, *p* = 0.13; experiment × condition × trial type: *F*_0.69,38_ = 0.27, *p* = 0.61). Therefore, these results replicate Experiment 1 results and add support for a strict hierarchical relationship between the sequence and task levels.

**Table 8.**
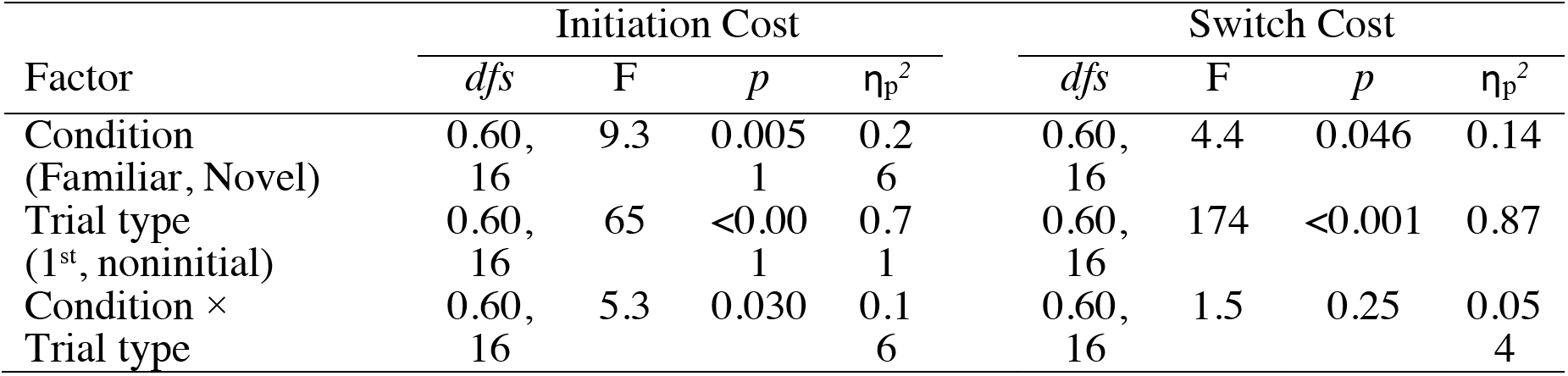
Experiment 2 rmANOVA for RT initiation cost (left) and switch cost (right)

**Table 9.**
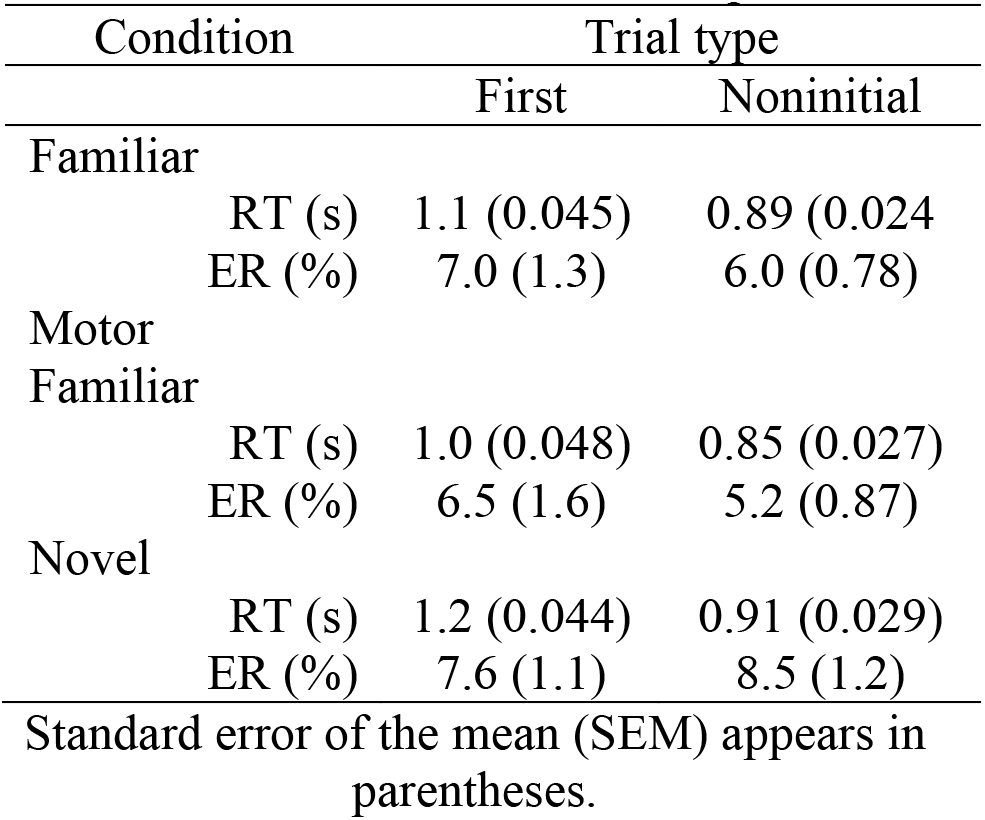
Condition and Trial type values for Reaction Time and Error Rate in Experiment 2

A strict relationship between the sequence and task levels is also supported by results from examining switch costs. While the increased practice in Experiment 2 did produce slightly faster RTs in Familiar sequences (*F*_0.50,14_ = 4.4, *p* = 0.046, η_p_^2^= 0.14; **Figure 5D**;**Table 8**), these effects were not specific to switch costs (i.e., there was no interaction) and were not different from Experiment 1 (condition [Familiar, Novel] × trial type [switch, repeat] × experiment [Experiment 1, Experiment 2] rmANOVA: experiment: *F*_0.55,31_ = 0.39, *p* = 0.53; experiment × condition: *F*_0.55,31_ = 2.00, *p* = 0.16; experiment × trial type: *F*_0.55,31_ = 2.7, *p* = 0.10; experiment × condition × trial type: *F*_0.55,31_ = 0.38, *p* = 0.65). Further, as in Experiment 1, additional foreknowledge did not modulate the relationship between switching and repeating at the first or subsequent positions (*F*_0.49,0.49,13_ = 0.75, *p* = 0.39, η_p_^*2*^= 0.027; **Table 10**). These results suggest that with additional practice the task level does not influence the sequence level, supporting a strict hierarchical relationship.

**Table 10.** Experiment 2 three-way rmANOVA with condition, trial type and switch type factors for RT.

| Factor | Reaction Time |  |  |  |
| --- | --- | --- | --- | --- |
| | *dfs* | F | *p* | $\eta_p^2$ |
| Condition | 0.49, | 8.9 | 0.0059 | 0.24 |
| (Familiar, Novel) | 13 |  |  |  |
| Trial type | 0.49, | 67 | <0.001 | 0.71 |
| (1<sup>st</sup>, noninitial) | 13 |  |  |  |
| Switch type | 0.49, | 73 | <0.001 | 0.73 |
| (sw, rep) | 13 |  |  |  |
| Condition × | 0.49, | 5.0 | 0.034 | 0.16 |
| Trial type | 13 |  |  |  |
| Condition × | 0.49, | 3.6 | 0.07 | 0.12 |
| Switch type | 13 |  |  |  |
| Trial type ×<br>Switch type | 0.49,<br>13 | 86 | <0.001 | 0.76 |
| Condition ×<br>Trial type ×<br>Switch type | 0.49,<br>0.49,<br>13 | 0.75 | 0.39 | 0.027 |

To examine the second question in Experiment 2, whether the addition of a motor sequence affected the relationship among the hierarchical levels, we performed planned comparisons between Motor Familiar and Novel sequences, and Motor Familiar and Familiar sequences. If there is a strict hierarchical relationship between sequence and motor level information, the addition of the motor sequence should not cause specific reductions in initiation costs. In contrast, a non-strict hierarchical relationship would allow for reductions in initiation costs due to motor-level sequential foreknowledge. First, we examined the relationship between the motor and abstract task sequence levels. Participants were faster (condition: *F*_0.68,18_ = 24, *p* < 0.001, η_p_^2^= 0.47) and initiation costs reduced (interaction: *F*_0.68,18_ = 12, *p* = 0.0016, η_p_^2^= 0.31) in Motor Familiar sequences compared to Novel sequences (**Figure 5C**). However, we cannot ascribe these improvements specifically to the addition of the motor sequence because Motor Familiar sequences contained practiced abstract task sequences, and we have also shown that practice improved performance on Familiar sequences. Therefore, we isolated the relationship between the motor and abstract task sequence levels by comparing Motor Familiar and Familiar sequence initiation costs. While Motor Familiar sequences were faster than Familiar sequences, these effects were not specific to initiation costs (interaction: *F*_0.75,20_ = 1.3, *p* = 0.26, η_p_^2^= 0.046, **Figure 5C**; **Table 11**). These results suggest that foreknowledge at the motor level does not affect processing at the sequence level and thus demonstrates a strict hierarchical relationship.

**Table 11.** Experiment 2 rmANOVA for RT initiation cost (left) and switch cost (right).

| Factor | Initiation Cost |  |  |  | Switch Cost |  |  |  |
| --- | --- | --- | --- | --- | --- | --- | --- | --- |
| | *dfs* | F | *p* | $\eta_p^2$ | *dfs* | F | *p* | $\eta_p^2$ |
| Condition<br>(Familiar, Motor<br>Familiar) | 0.75,<br>20 | 7.8 | 0.009<br>5 | 0.22 | 0.60,<br>16 | 5.7 | 0.024 | 0.18 |
| Trial type<br>(1<sup>st</sup>, noninitial) | 0.75,<br>20 | 42 | <0.00<br>1 | 0.61 | 0.60,<br>16 | 123 | <0.001 | 0.82 |
| Condition ×<br>Trial type | 0.75,<br>20 | 1.3 | 0.26 | 0.04<br>6 | 0.60,<br>16 | 18 | <0.001 | 0.39 |

In contrast, we found support for a non-strict hierarchical relationship between the task and motor levels. Participants exhibited reduced task-switch costs in Motor Familiar sequences compared to Familiar sequences (interaction: *F*_0.6,16_ = 17, *p* < 0.001, η_p_^2^= 0.39, **Figure 5D**;**Table 11**) that were driven by reductions in switch trial RTs (post-hoc t-test, Bonferroni-adjusted a = 0.025: repeat trials, *t*_27_ = 0.16, *p* = 0.87, *d* = 0.031; switch trials, *t*_27_ = 3.8, *p* < 0.001, *d* = 0.40). Further, switching and repeating at the motor level interacted with switching and repeating at the task level. In this congruency effect, task switches accompanied by motor repeats were executed more slowly than task switches accompanied by motor switches (motor response type [repeat, switch] × trial type [repeat, switch] rmANOVA: motor response type × task trial type: *F*_0.47,13_ = 96, *p* < 0.001, η_p_^*2*^= 0.78; **Table 12**). This result replicates previous studies that were not performed in the context of abstract task sequences (Kikumoto & Mayr, 2020; Korb et al., 2017; Mayr & Bryck, 2005). Together, these results suggest that the motor level influences the task level and support a non-strict hierarchical relationship between them.

**Table 12.**
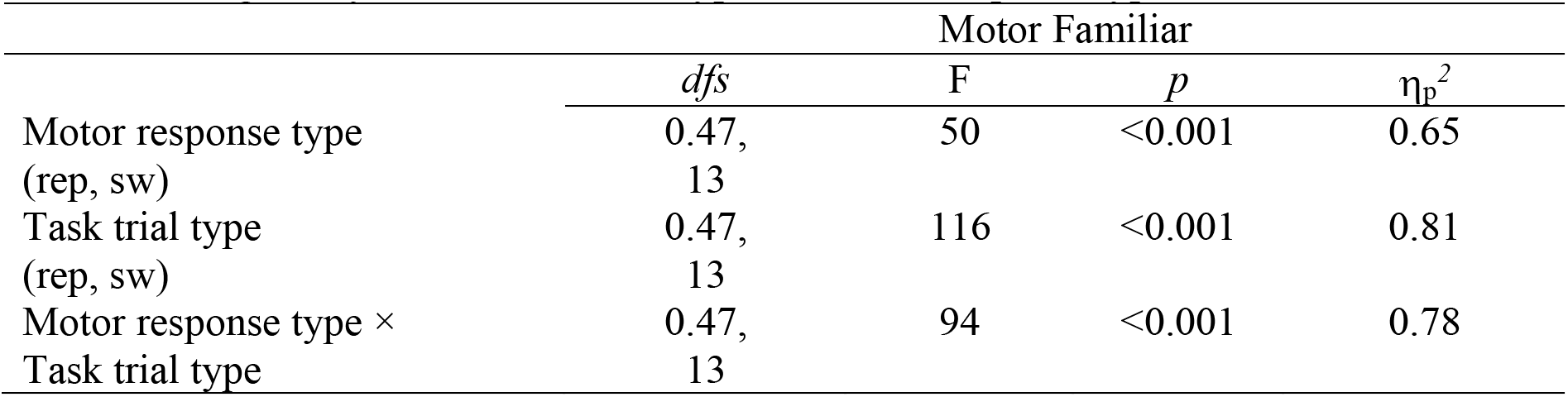
Congruency between task trial type and motor response type rmANOVA on RT.

|  | Motor Familiar |  |  |  |
| --- | --- | --- | --- | --- |
| | *dfs* | F | *p* | $\eta_p^2$ |
| Motor response type<br>(rep, sw) | 0.47,<br>13 | 50 | <0.001 | 0.65 |
| Task trial type<br>(rep, sw) | 0.47,<br>13 | 116 | <0.001 | 0.81 |
| Motor response type ×<br>Task trial type | 0.47,<br>13 | 94 | <0.001 | 0.78 |

Two control analyses support the isolable influence of the motor level on the task level. First, a possible design concern is that Motor Familiar sequences necessarily had one button press that occurred more frequently than other responses, due to the presence of a motor repeat trial in the motor sequence. In contrast, button press response frequency was balanced in Familiar and Novel blocks of trials. We mitigated this aspect of the task design with counterbalancing, as each participant learned two different motor sequences (see Methods). Additionally, we performed follow-up analyses to verify that our effects were likely not caused by response frequency effects. It is possible that participants were faster at the most frequent response finger if frequency effects were driving RT reductions. We tested this possibility and found that participants were not faster at more frequent finger responses in Motor Familiar compared to Familiar blocks (*F*_0.45,12_ = 2.2, *p* = 0.15, η_p_^*2*^= 0.076; **Table 13**), indicating that responding more frequently with one finger did not yield RT reductions. Further, if these effects were due solely to finger frequency rather than sequential content, the effect would be uniform across response repeats and switches Thus, we assessed response switching and repeating across conditions. We found that there were greater response switch costs in Motor Familiar compared to Familiar blocks (motor response type [motor repeat, motor switch] × condition [Motor Familiar, Familiar] rmANOVA: *F*_0.45,12_ = 6.9, *p* = 0.014, η_p_^*2*^= 0.20; **Table 13**), suggesting that these differences were not due to frequency effects alone.

**Table 13.**
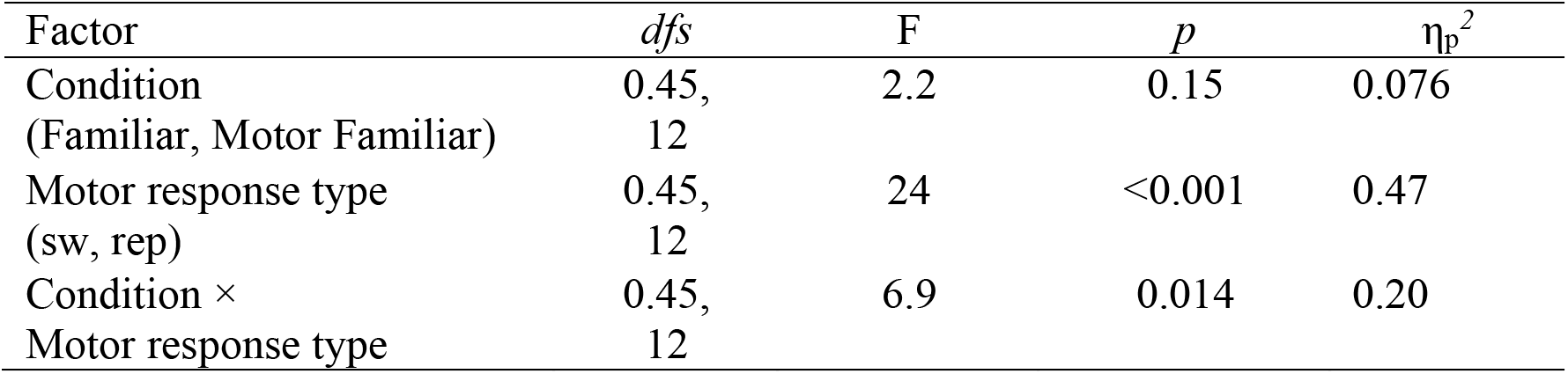
Two-way rmANOVA including trials where the correct response was the most frequent finger for RT.

Second, awareness of the embedded motor sequences may influence their impact on superordinate levels in the hierarchy, as awareness improves performance on practiced motor sequences (Curran & Keele, 1993). We did not find evidence that participants were aware of embedded motor sequences. First, none of the participants reproduced the motor sequence they used during the task (*n* = 28); however, on the post-test questionnaire some did report awareness of a repeating pattern (*n* = 7). Second, participants were significantly slower when executing the Motor Familiar sequences (*M =* 0.91 s, *SD* = 0.17 s) compared to the motor sequences in the SRTT after the test phase (*M =* 0.42 s, *SD* = 0.091 s; *t*_27_ = 15, *p* < 0.001, *d* = 3.7), suggesting that participants continued to execute the tasks throughout the experimental session and did not shift to executing the motor sequences. Third, in response to the question at the end of each block to guess the next correct key press (without a stimulus present), participants’ performance was not different from chance (chance = 33%; *M* = 35%, *SD* = 9.2%, *t*_27_ = 0.77, *p* = 0.45, *d* = 0.21). Taken together, these three assessments indicated that participants were not explicitly aware of the embedded motor sequences and continued to execute the abstract task sequences in the Motor Familiar condition.

In summary, results from Experiment 2 provide evidence that additional practice does not further reduce initiation costs or alter the apparently strict hierarchical relationship between the sequence and task levels. Further, the addition of an embedded motor sequence improved abstract task sequence performance in an additive manner, suggesting a strict hierarchical relationship between the sequence and motor levels as well. In contrast, we provide evidence for a non-strict hierarchical relationship between task-level and motor-level information. Because this relationship is evident in the Motor Familiar condition where the embedded motor sequence and tasks are practiced together, it is possible that the influence of the motor level on the task level could be due to an integrated representation (Cock & Meier, 2013; Mayr & Bryck, 2005; Weiermann et al., 2010). Therefore, an open question is whether the non-strict interaction between the motor and task levels is due to the joint practice, or whether they can be observed separately. In Experiment 3, we will replicate Experiment 2 and address this question.

## Experiment 3

The objective of Experiment 3 was to further examine interactions between the motor level and superordinate levels. We tested whether the interdependencies were emergent with practice or inherent to the structure of the hierarchical task, and whether participants formed an integrated representation of the abstract sequence and motor sequence information. We replicated the experimental design of Experiment 2 with a separate set of participants and added a probe phase after the test phase. During the probe phase, we removed embedded motor sequences from the Motor Familiar condition, making it more similar to the Familiar condition during the test phase. We also added embedded motor sequences to the Familiar condition in the probe phase, making it more similar to the Motor Familiar condition during the test phase. We hypothesized that abstract task and motor sequences practiced together would not form an integrated representation of task elements and that interdependencies between hierarchical levels were inherent to the execution of the task.

## Methods

### Participants

Recruitment, inclusion criteria, and consenting procedure were the same as in Experiments 1 and 2. Thirty-three participants participated in the study. Participants with over 20% error were excluded (*n* = 6). Twenty-seven (*n* = 17 female) adults between the ages of 18-35 (*M =* 21 *SD* = 2.9) were included in this study.

### Procedure

Probe phase: The procedure in Experiment 3 was identical to the procedure in Experiment 2 except for the addition of the probe phase. The probe phase included three runs after the completion of the six test runs during the second session. The structure of these runs was identical to the test runs (8 blocks each). However, the blocks were composed of two new trial types: Motor Familiar Minus and Familiar Plus. The Motor Familiar Minus condition consisted of the Motor Familiar abstract sequences that participants had practiced (one simple and one complex) with the four-member embedded motor sequences removed. Instead, correct button press responses were randomized such that they followed no predictable motor sequence. The Familiar Plus condition included the Familiar abstract sequences that participants had practiced (one simple and one complex) but now with embedded motor response sequences originally learned as part of the Motor Familiar sequences. The motor sequences transferred from the complex Motor Familiar condition to the complex Familiar Plus condition, and from the simple Motor Familiar condition to the simple Familiar Plus condition. Each probe run consisted of two blocks of each sequence condition. Participants were not instructed that these runs would be different in any way.

After the completion of the post-task questionnaire, participants completed the SRTT as in Experiment 2. During this version of the SRTT, participants completed two blocks of trials with one predictable motor sequence, one random block. Another two blocks of trials with the other predictable motor sequence, and a final random block.

### Analysis

Statistical analyses were conducted in Matlab (MathWorks; RRID:SCR_001622). RTs and ERs were submitted to rmANOVAs and t-tests where appropriate. To examine the test phase of Experiment 3 that replicated Experiment 2, we performed the same planned condition × trial type rmANOVA comparisons between Familiar and Novel conditions, Motor Familiar and Familiar conditions. We conducted analyses addressing motor response frequency, congruency between the motor and task level, and the interaction between switching and repeating at the first position as in Experiment 2. We also conducted across experiment analyses by including group (Experiment 2 or 3) as a factor in the rmANOVA analyses. We further compared across experiments by including awareness (aware, unaware) during the test phase as a factor in the rmANOVA analyses. To examine the probe phase we performed planned comparisons in the same way (condition × trial type rmANOVAs) between probe conditions (Motor Familiar Minus, Familiar Plus), and between conditions that contained the same abstract sequence, but differed between probe and test as to whether or not they contained an embedded motor sequence (Familiar, Familiar Plus and Motor Familiar, Motor Familiar Minus). Additionally, we compared performance on probe conditions to performance on Novel sequences (Motor Familiar Minus, Novel). As in the previous experiments, our primary comparisons were concerned with differences in RT between conditions while comparing average RT of switch trials to average RT of repeat trials or comparing average RT at the first position compared to the unweighted average RT of noninitial positions (see Experiment 1 Methods). We again combined across complexity conditions. When sphericity assumptions were violated, we used Greenhouse Geisser correction for the degrees of freedom.

## Results

The test phase of Experiments 2 and 3 were identical. We first replicated analyses from Experiment 2, and then examined the probe phase of Experiment 3 to determine if abstract and motor sequences formed an integrated representation.

Participant performance in the test phase of Experiment 3 was similar to performance in Experiment 2. Overall ER at test was 6.1% (*SD* = 3.7%). Again, we assessed the learning of Familiar and Motor Familiar sequences first by comparing performance during the practice phase to performance at test. RTs were faster in Familiar and Motor Familiar sequences at test as compared practice (Familiar: *t*_26_ = 12, *p* < 0.001, *d* = 1.2; Motor Familiar: *t*_26_ = 7.5, *p* < 0.001, *d* =1.3; **Figure 6A**). ER was marginally lower at test for Familiar sequences (ER: Familiar: *t*_26_ =2.0, *p* = 0.058, *d* = 0.40; **Figure 6B**) and significantly lower in Motor Familiar sequences (ER: Motor Familiar: *t*_26_ =3.2, *p* = 0.0036, *d* = 0.51; **Figure 6B**). Initiation costs were observed in RT and not in ER (**Figure 6C**; **Table 14**). All subsequent analyses combined across simple and complex sequences, as there was no difference between them (*F*_0.68,18_ = 0.28, *p* = 0.60, η_p_^2^= 0.011), and primarily address RT performance in the test and probe phases.

**Figure 6.**
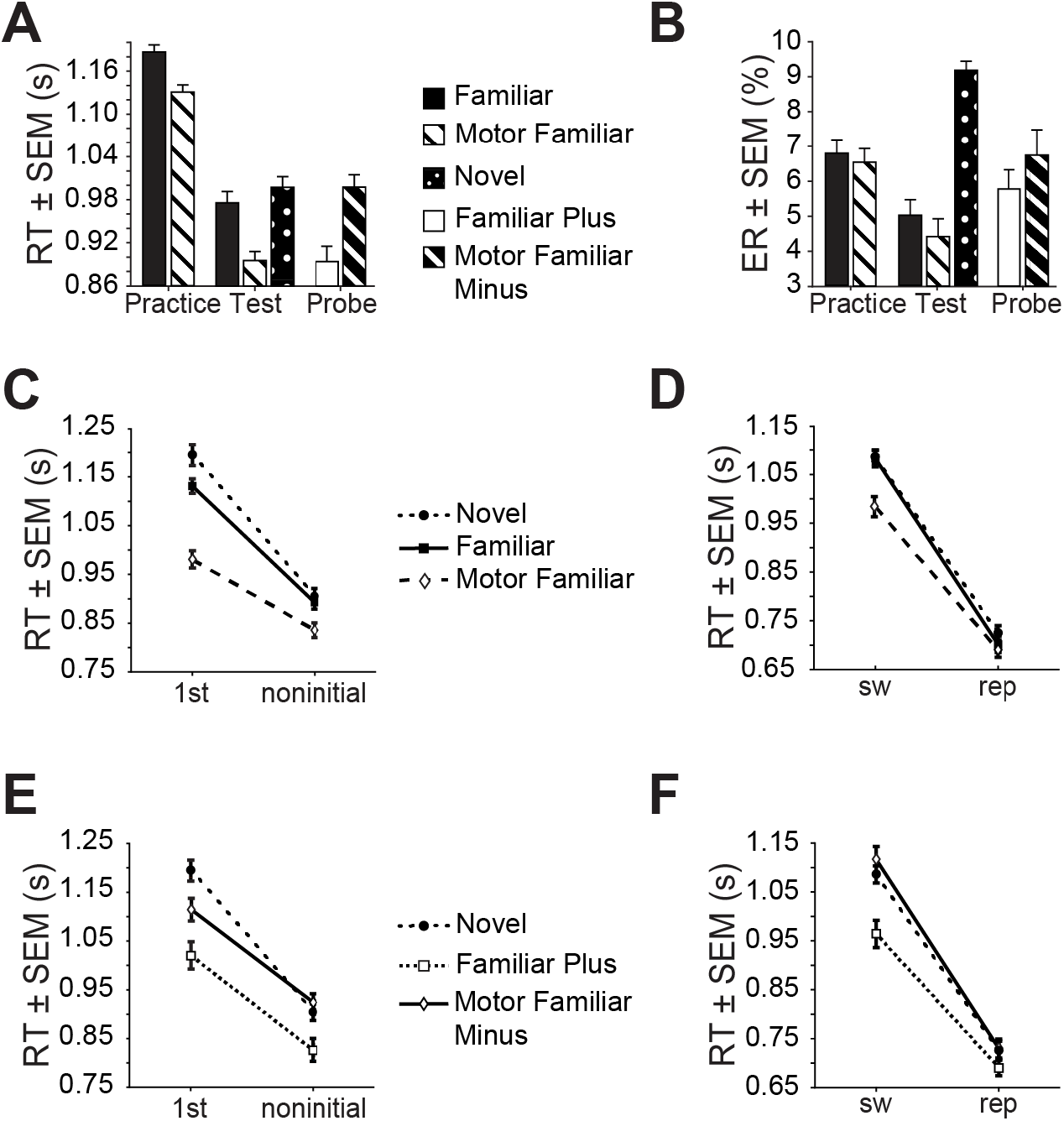
Experiment 3 Results. (**A**) Average reaction time (ERTR) plotted by study phase and condition. (**B**) Average error rate (ER) as in A. (**C**) Plot of RT for 1^st^ versus noninitial trial types by condition during the test phase. **(D)** Plot of RT for switch (sw) versus repeat (rep) trial types by condition during the test phase. **(E)** Plot of RT for 1^st^ versus noninitial trial types by condition during the probe phase. **(F)** Plot of RT for switch (sw) versus repeat (rep) trial types by condition during the probe phase. Familiar/solid/squares; Motor Familiar/dashed/diamonds; Novel/dotted/circles; Familiar Plus/dashed/squares; Motor Familiar Minus/solid/diamonds.

**Table 14.** Experiment 3 t-tests comparing first and noninitial positions for RT and ER.

| Condition | dfs | Reaction Time |  |  | Error Rate |  |  |
| --- | --- | --- | --- | --- | --- | --- | --- |
|  |  | t | p | d | t | p | d |
| Familiar | 26 | 9.9 | <0.001 | 1.3 | -0.29 | 0.78 | 0.046 |
| Motor Familiar | 26 | 5.9 | <0.001 | 0.81 | 0.33 | 0.75 | 0.046 |
| Novel | 26 | 12 | <0.001 | 1.5 | -2.4 | 0.022 | 0.342 |

Reaction Time and Error Rate in Experiment 3The effects of sequence level practice and support for a strict hierarchical structure between the task and sequence levels were consistent across experiments. Initiation costs were specifically reduced in Familiar compared to Novel sequences (interaction:*F*_0.65,17_ = 5.5, *p* = 0.027, η_p_^2^= 0.17; post-hoc t-test, Bonferroni-adjusted a = 0.025: first positions, *t*_26_ = −3.0, *p* = 0.0063, *d* = 0.29; noninitial, *t*_26_ =-1.3, *p* = 0.20, *d* = 0.095; **Figure 6C**; **Table 15** and **Table 16**). Further, task-level switch costs were not altered with practice (interaction: *F*_0.53,14_ = 1.2, *p* = 0.28, η_p_^2^= 0.044; **Figure 6D**; **Table 15**), and the relationship between switching and repeating was not altered at the first or subsequent positions (condition × trial type × switch type: *F*_0.49,0.49,13_ = 2.6, *p* = 0.12, η_p_^*2*^= 0.092; **Table 17**). Together, these results replicate Experiments 1 and 2 and support a strict hierarchical model of abstract task sequence execution between the sequence and task levels.

**Table 15.** Experiment 3 rmANOVA for RT initiation cost (left) and switch cost (right).

| Factor | Initiation Cost |  |  |  | Switch Cost |  |  |  |
| --- | --- | --- | --- | --- | --- | --- | --- | --- |
| | dfs | F | p | $\eta_p^2$ | dfs | F | p | $\eta_p^2$ |
| Condition (Familiar, Novel) | 0.65, 17 | 9.2 | 0.0054 | 0.26 | 0.53, 13.8 | 1.7 | 0.20 | 0.063 |
| Trial type (1<sup>st</sup>, noninitial) | 0.65, 17 | 132 | <0.001 | 0.84 | 0.53, 13.8 | 221 | <0.001 | 0.89 |
| Condition × Trial type | 0.65, 17 | 5.5 | 0.027 | 0.17 | 0.53, 13.8 | 1.2 | 0.28 | 0.044 |

**Table 16.**
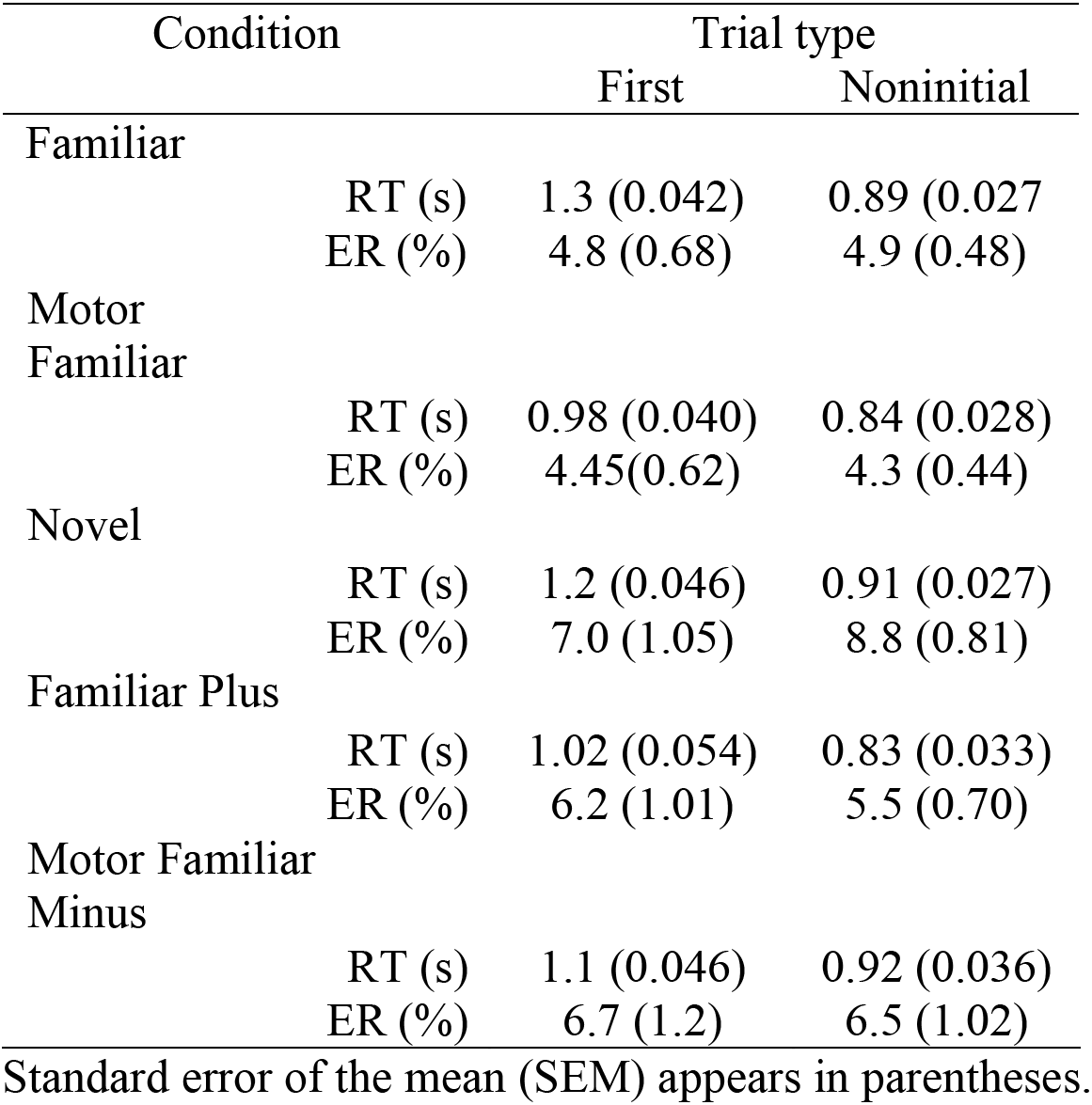
Condition and Trial type values for Reaction Time and Error Rate in Experiment 3

**Table 17.** Experiment 3 three-way rmANOVA with condition, trial type and switch type factors for RT.

| Factor | Reaction Time |  |  |  |
| --- | --- | --- | --- | --- |
| | *dfs* | F | *p* | $\eta_p^2$ |
| Condition<br>(Familiar, Novel) | 0.49,<br>13 | 8.5 | 0.007 | 0.25 |
| Trial type<br>(1<sup>st</sup>, noninitial) | 0.49,<br>13 | 130 | <0.001 | 0.83 |
| Switch type<br>(sw, rep) | 0.49,<br>13 | 124 | <0.001 | 0.83 |
| Condition ×<br>Trial type | 0.49,<br>13 | 5.2 | 0.031 | 0.17 |
| Condition ×<br>Switch type | 0.49,<br>13 | 5.3 | 0.030 | 0.17 |
| Trial type ×<br>Switch type | 0.49,<br>13 | 50 | <0.001 | 0.66 |
| Condition ×<br>Trial type ×<br>Switch type | 0.49,<br>0.49,<br>13 | 2.6 | 0.12 | 0.092 |

To isolate the effects of embedded motor sequences and assess the relationship between the motor and sequence levels, we compared Motor Familiar sequences to Familiar sequences in the Experiment 3 test phase. The relationship between the motor and abstract task sequence levels in the test phase of Experiment 3 was similar to the relationship observed in Experiment 2. Again, RTs on Motor Familiar sequences were faster than Familiar sequences (condition: *F*_0.75,20_ = 27, *p* < 0.001, η_p_^2^=0.51; **Table 18**). However, in potential contrast to Experiment 2, in Experiment 3 initiation costs were specifically reduced in Motor Familiar sequences (interaction: *F*_0.75,20_ = 28, *p* < 0.001, η_p_^2^= 0.52; **Figure 6C** and **Table 18**; post-hoc t-test, Bonferroni-adjusted a = 0.025: first position, *t*_26_ = 6.6, *p* < 0.001, *d* = 0.71; noninitial positions, *t*_26_ = 2.7, *p* = 0.013, *d* = 0.39). These results may indicate that practice with embedded motor sequences was different between Experiments 2 and 3. To investigate this possibility, we compared the relationship between Familiar and Motor Familiar sequences across the two experiments. We found no reliable differences between the experiments (condition [Familiar, Motor Familiar] × trial type [first, noninitial] × experiment [Experiment 2, Experiment 3] rmANOVA, experiment: *F*_0.78,41_ =0.021, *p* = 0.88; condition × experiment: *F*_0.78,41_ = 1.8, *p* = 0.19; trial type × experiment: *F*_0.78,41_ = 0.0024, *p* = 0.96; condition × trial type × experiment: *F*_0.78,41_ = 1.65, *p* = 0.20). Therefore, though there appeared to be differences in the effects of adding an embedded motor sequence beyond that of sequence practice alone between Experiments 2 and 3, these differences were not reliable and the results in Experiment 3 primarily replicated the results in Experiment 2.

**Table 18.** Experiment 3 rmANOVA for RT initiation cost (left) and switch cost (right).

| Factor | Initiation Cost |  |  |  | Switch Cost |  |  |  |
| --- | --- | --- | --- | --- | --- | --- | --- | --- |
| | *dfs* | F | *p* | $\eta_p^2$ | *dfs* | F | *p* | $\eta_p^2$ |
| Condition<br>(Familiar, Motor<br>Familiar) | 0.75,<br>20 | 27 | <0.00<br>1 | 0.51 | 0.70,<br>18 | 7.2 | 0.013 | 0.22 |
| Trial type<br>(1<sup>st</sup>, noninitial) | 0.75,<br>20 | 72 | <0.00<br>1 | 0.73 | 0.70,<br>18 | 172 | <0.001 | 0.87 |
| Condition ×<br>Trial type | 0.75,<br>20 | 28 | <0.00<br>1 | 0.52 | 0.70,<br>18 | 22 | <0.001 | 0.46 |

At the level of task and motor interactions, results supported a non-strict hierarchical relationship between the task and motor levels as in Experiment 2. Switch costs were selectively reduced in Motor Familiar sequences (interaction: *F*_0.7,18_ = 22, *p* < 0.001, η_p_^2^= 0.46; post-hoc t-test, Bonferroni-adjusted a = 0.025: repeat trials: *t*_26_ = 0.64, *p* = 0.53, *d* = 0.10; switch trials: *t*_26_ =3.7, *p* < 0.001, *d* = 0.51, **Figure 6D**; **Table 18**). Motor response switching and repeating also interacted with task switching and repeating such that task switches with response repeats were slower than those with response switches (motor response type [repeat, switch] × task trial type [repeat, switch]: *F*_0.52,13_ = 137, *p* <0.001, η_p_^2^= 0.84), replicating Experiment 2 and previous work (Kikumoto & Mayr, 2020; Korb et al., 2017; Mayr & Bryck, 2005). Together, these results provide evidence of the influence of the motor level on the task level and suggest a non-strict hierarchical relationship.

As in Experiment 2, we performed two additional sets of control analyses to isolate the effects of embedded motor sequences from potential effects of response frequency and awareness. We again found that responses with the more frequent finger (used for repeat trials) were not faster in Motor Familiar blocks as compared to Familiar blocks (*F*_0.52,14_ = 1.8, *p* = 0.19, η_p_^2^= 0.065) and that there were greater switch costs in Motor Familiar blocks as compared to Familiar blocks (*F*_0.52,14_ = 5.2, *p* = 0.031, η_p_^2^= 0.17; **Table 19**), indicating that finger frequency alone was not a primary driver of the reductions in RT in the Motor Familiar blocks. Second, to address awareness of the motor sequences, no participants accurately reproduced the motor sequences (N = 27), although 12 participants reported noticing a repeating pattern in the responses for some sequences. This recognition did not affect other measures of awareness that we examined. Participants performed at chance on the button guess judgement (33%; *M* = 34 %, *SD* = 12%, *t*_26_ = 0.34, *p* = 0.74, *d* = 0.095), and were faster on the SRTT alone (Motor Familiar: *M =* 0.90 ± 0.16; *SRTT: M =* 0.46 ± 0.089 s; *t*_26_ = 16, *p* < 0.001, *d* = 3.4). Therefore, while participants’ awareness was limited, it is possible that the heightened recognition of patterns in the motor responses may have contributed to additional improvements in control costs at the sequence level as evidenced by reductions at the first position of abstract task sequences.

**Table 19.** Two-way rmANOVA including trials where the correct response was the most frequent finger for RT.

| Factor | Reaction Time |  |  |  |
| --- | --- | --- | --- | --- |
| | *dfs* | F | *p* | $\eta_p^2$ |
| Condition<br>(Familiar, Novel) | 0.52,<br>14 | 1.8 | 0.19 | 0.065 |
| Trial type<br>(motor: sw, rep) | 0.52,<br>14 | 20 | <0.001 | 0.43 |
| Condition ×<br>Trial type | 0.52,<br>14 | 5.2 | 0.031 | 0.17 |

To address the hypothesis that participant awareness could influence the potential for the motor-level to influence the abstract task sequence-level, and compensate for the small number of participants that may have been aware of the motor sequences in each experiment, we combined Experiments 2 and 3 to compare aware and unaware participants. We did not find evidence that awareness modulated the relationship between initiation costs for Motor Familiar and Familiar conditions (Awareness × Condition × Trial type: *F*_0.76,0.76,40_ = 0.57, *p* =0.45, **Table 20**). However, there was statistical evidence that awareness differentially affected Motor Familiar sequences compared to Familiar sequences (Awareness × Condition: *F*_0.76,40_ = 6.8, *p* =0.012, **Table 20**). Because there were no embedded motor sequences in the Familiar condition to become aware of, this result suggests that changes in the performance of Motor Familiar sequences due to awareness may drive this effect. Together these results can neither support nor rule out the hypothesis that awareness could influence the relationship between the motor and sequence levels. Further experiments will be necessary to address this question.

**Table 20.** Three-way rmANOVA on RT data with Awareness (aware, unaware) and Experiment (Experiment 2, Experiment 3) as between group factors and Condition (familiar, novel) and Trial type (first, noninitial) as within group factors.

| Factor | Reaction Time |  |  |  |
| --- | --- | --- | --- | --- |
| | *df* | *F* | *p* | $\eta_p^2$ |
| Experiment<br>(Exp 2, Exp 3) | 0.76,<br>40 | 0.035 | 0.85 |  |
| Awareness<br>(aware, unaware) | 0.76,<br>40 | 2.7 | 0.11 |  |
| Condition<br>(Familiar, Novel) | 0.76,<br>40 | 40 | <0.001 | 0.44 |
| Trial type<br>(1<sup>st</sup>, noninitial) | 0.76,<br>40 | 88 | <0.001 | 0.63 |
| Awareness ×<br>Condition | 0.76,<br>40 | 6.8 | 0.012 |  |
| Experiment ×<br>Condition | 0.76,<br>40 | 0.70 | 0.41 |  |
| Awareness ×<br>Trial type | 0.76,<br>40 | 1.2 | 0.28 |  |
| Experiment ×<br>Trial type | 0.76,<br>40 | 0.031 | 0.86 |  |
| Condition ×<br>Trial type | 0.76,<br>40 | 8.4 | 0.0054 | 0.14 |
| Awareness ×<br>Condition ×<br>Trial type | 0.76,<br>0.76,<br>40 | 0.57 | 0.45 |  |
| Experiment ×<br>Condition ×<br>Trial type | 0.76,<br>0.76,<br>40 | 2.0 | 0.17 |  |

To examine whether practicing an abstract sequence with embedded motor sequences may facilitate performance through the creation of an integrated representation, we examined the probe phase in Experiment 3 in a series of planned comparisons. In the probe phase, we removed the embedded motor sequences from the test phase Motor Familiar abstract task sequences to create the Motor Familiar Minus probe condition, and we added those same motor sequences to the test phase Familiar abstract task sequence to create the Familiar Plus probe condition. Results from this manipulation are presented below.

Results from Experiments 1 and 2 and the test phase of Experiment 3 support the hypothesis that there is a strict hierarchical relationship between the sequence and task levels. In this context, we hypothesized that the relationship between the sequence and task levels would not be altered by the probe block manipulation, as effects would be primarily dictated by abstract task sequences. In the test phase we examined this relationship by comparing Familiar and Novel sequences. In the probe phase, the analogous comparison is between abstract task sequences without embedded motor sequences, Motor Familiar Minus, and Novel sequences. We found a decrease in RT selectively at the first position in the sequence (condition [Novel, Motor Familiar Minus] × trial type [first, noninitial] rmANOVA, condition: *F*_0.76,20_ = 2.1, *p* = 0.16, η_p_^2^= 0.074, interaction: *F*_0.76,20_ = 10, *p* = 0.0036, η_p_^2^= 0.28, **Figure 6E**, **Table 21**; post-hoc t-test, Bonferroni-adjusted a = 0.025, first position, *t*_26_ = 2.6, *p* = 0.015, *d* = 0.035; noninitial positions, *t*_26_ = 0.77, *p* = 0.45, *d* = 0.11). This result supports a strict hierarchical relationship between the sequence and task levels that is not integrated during practice.

**Table 21.** Experiment 3 probe phase rmANOVA on RT initiation cost.

| Factor | Initiation Cost |  |  |  |
| --- | --- | --- | --- | --- |
| | *dfs* | F | *p* | $\eta_p^2$ |
| Condition<br>(Novel,<br>Motor Familiar Minus) | 0.76,<br>20 | 2.1 | 0.16 | 0.074 |
| Trial type<br>(1<sup>st</sup>, noninitial) | 0.76,<br>20 | 104 | <0.001 | 0.80 |
| Condition ×<br>Trial type | 0.76,<br>20 | 10 | 0.0036 | 0.28 |

Results from Experiments 2 and 3 test blocks generally supported a strict hierarchical relationship between the abstract task sequence and motor levels, though there may be other factors that influence that relationship (addressed further in the Discussion). We hypothesized that the probe block manipulation would not alter the relationship between the motor and abstract task sequence levels if there was a strict hierarchical relationship and, therefore, no potential for integration. We first showed that the probe block conditions were not reliably different from their counterpart test block conditions. Specifically, the abstract task sequences without embedded motor sequences in test were not reliably different from probe (condition [Familiar, Motor Familiar Minus] × trial type [first, noninitial] rmANOVA, condition: *F*_0.70,18_ = 0.82, *p* =0.78, η_p_^2^= 0.0031; trial type: *F*_0.70,18_ = 86.4, *p* <0.001, η_p_^2^= 0.77; interaction: *F*_0.70,18_ = 3.0, *p* =0.097, η_p_^2^= 0.10; **Figure 6C** and **Figure 6E**). The same was true for abstract task sequences with embedded motor sequences during the test and probe phases (condition [Motor Familiar, Familiar Plus] × trial type [first, noninitial] rmANOVA, condition: *F*_0.62,16_ = 0.47, *p* = 0.50, η_p_^2^= 0.018; trial type: *F*_0.62,16_ = 53 *p* <0.001, η_p_^2^= 0.67; interaction: *F*_0.62,16_ = 3.6, *p* = 0.069, η_p_^2^= 0.12; **Figure 6C** and **Figure 6E**). These results suggest that any differences in the relationship between the probe conditions could not be due to a failure to transfer the embedded motor sequences.

To specifically address the question of integration within the hierarchy between the sequence and motor levels, we examined initiation costs in the probe block conditions: Motor Familiar Minus and Familiar Plus. We hypothesized that if participants had formed an integrated representation, then the Motor Familiar Minus condition would show a decrement in performance beyond the subtraction of the embedded motor sequence and the Familiar Plus condition would not show the same facilitation as its test phase counterpart, the Motor Familiar condition. In contrast, if participants were not forming an integrated representation then we hypothesized that the performance of the Motor Familiar Minus and Familiar Plus conditions would be comparable to their test phase counterparts, Familiar and Motor Familiar respectively. In contrast to the test phase of Experiment 3, the addition of the embedded motor sequence in the probe phase did not selectively affect initiation costs (interaction: *F*_0.74,19_ = 0.0098, *p* = 0.92, η_p_^2^= 0.00038, **Figure 6E**, **Table 22**). This effect is the same as in Experiment 2, where a strict hierarchical relationship between the sequence and motor levels was observed without an interaction specific to initiation costs (condition [with or without embedded motor] × trial type [first, noninitial] × experiment [Experiment 2, Experiment 3], rmANOVA, condition × trial type × experiment interaction: *F*_0.8,42_ = 0.94, *p* = 0.33). Moreover, this result contrasted with the pattern of results in the test phase of Experiment 3 (condition [with or without embedded motor] × phase [test, probe] × trial type [first, noninitial], rmANOVA, condition × phase × trial type: *F*_0.80,42_ =5.84, *p* =0.02, η_p_^2^= 0.19). Together, these results suggest that the relationship between the abstract task sequence and motor levels is strict, without integration, and the effects of an embedded motor sequence are additive in the speeding of abstract task sequence execution. The apparent differences between Experiment 3 test and probe and the potential effects of practice on these relationships will be explored further in the Discussion.

**Table 22.** Experiment 3 probe phase rmANOVA on RT initiation cost (left) and switch cost (right).

| Factor | Initiation Cost |  |  |  | Switch Cost |  |  |  |
| --- | --- | --- | --- | --- | --- | --- | --- | --- |
| | *dfs* | F | *p* | $\eta_p^2$ | *dfs* | F | *p* | $\eta_p^2$ |
| Condition<br>(Motor Familiar Minus, Familiar Plus) | 0.74,<br>19 | 9.6 | 0.0047 | 0.27 | 0.75,<br>19 | 10 | 0.003<br>9 | 0.28 |
| Trial type<br>(1<sup>st</sup>, noninitial) | 0.74,<br>19 | 54 | <0.001 | 0.67 | 0.75,<br>19 | 149 | <0.00<br>1 | 0.85 |
| Condition ×<br>Trial type | 0.74,<br>19 | 0.0098 | 0.92 | 0.00038 | 0.75,<br>19 | 11 | 0.002<br>6 | 0.30 |

In contrast to the relationship between the other hierarchical levels, we found evidence across Experiment 2 and 3 test blocks that there was a non-strict hierarchy between the motor and task levels, consistent with previous findings (Kikumoto & Mayr, 2020; Korb et al., 2017; Mayr & Bryck, 2005). Previous work suggests that there may be an integrated representation formed between stimuli and their responses when they are practiced. If there was an integrated representation formed during practice, then we hypothesized that the probe block manipulation would break this relationship and manifest in a lack of selective facilitation of the switch costs by the embedded motor sequence such that the effects would appear more additive. First, we confirmed that there were no overall differences between the conditions at test and probe on the task level (condition [Familiar, Motor Familiar Minus] × trial type [switch, repeat] rmANOVA, condition: *F*_0.72,19_ = 2.1, *p* = 0.16, η_p_^2^= 0.073; *F*_0.72,19_ = 168, *p* <0.001, η_p_^2^= 0.87; interaction: *F*_0.72,19_ = 0.043, *p* = 0.84, η_p_^2^= 0.0016; condition [Motor Familiar, Familiar Plus] × trial type [switch, repeat] rmANOVA, condition: *F*_0.60,16_ = 0.32, *p* = 0.57, η_p_^2^= 0.012; trial type: *F*_0.60,16_ = 121, *p* <0.001, η_p_^2^= 0.82; interaction: *F*_0.61,16_ = 1.8, *p* = 0.19, η_p_^2^= 0.065; **Figure 6D** and **Figure 6F**). To address the hierarchy question, we replicated the relationship we observed in the Experiment 2 and 3 test phases where RTs were reduced selectively for switch trials (interaction: *F*_0.75,20_ = 11, *p* = 0.0026, η_p_^2^= 0.30; **Figure 6F**, **Table 22**; post-hoc t-test, Bonferroni-adjusted a = 0.025: switch trials, *t*_26_ = 3.6, *p* = 0.0014, *d* = 0.64; repeat trials, *t*_26_ = 1.6, *p* = 0.12, *d* = 0.27). Further, we replicated the finding that switch costs at the motor and task-level interact in a nonadditive manner in the Motor Familiar Minus (motor response type [switch, repeat] × task trial type [switch, repeat]: *F*_0.59,15_ = 183, *p* < 0.001, η_p_^2^= 0.88) and Familiar Plus conditions (motor response type [switch, repeat] × task trial type [switch, repeat]: *F*_0.60,16_ = 105, *p* < 0.001, η_p_^2^= 0.80; **Table 23**). These results suggest that an integrated representation was not formed between the task and motor levels, despite the non-strict relationship between the levels. This finding has implications for understanding flexible behavior and interactions between motor and cognitive processing and will be explored further in the Discussion.

**Table 23.** Congruency between task trial type and motor response type rmANOVA on RT.

| Factor | Motor Familiar Minus |  |  |  | Familiar Plus |  |  |  |
| --- | --- | --- | --- | --- | --- | --- | --- | --- |
| | *dfs* | F | *p* | $\eta_p^2$ | *dfs* | F | *p* | $\eta_p^2$ |
| Motor response type<br>(repeat, switch) | 0.60,<br>16 | 29 | <0.00<br>1 | 0.27 | 0.59,<br>15 | 9.8 | <0.001 | 0.82 |
| Task trial type<br>(repeat, switch) | 0.60,<br>16 | 120 | <0.00<br>1 | 0.82 | 0.59,<br>15 | 119 | <0.001 | 0.53 |
| Motor response type ×<br>Task trial type | 0.60,<br>16 | 183 | <0.00<br>1 | 0.80 | 0.59,<br>15 | 105 | <0.001 | 0.88 |

## Discussion

These studies investigated the hierarchical relationships between levels of abstract task sequences. We manipulated practice at the sequence level and the presence of embedded response sequences at the motor level to operationalize foreknowledge and adjudicate between strict and non-strict hierarchical relationships at the sequence, task, and motor levels. There were four main findings across three experiments. First, we provided consistent evidence for a strict hierarchical relationship between sequence and task levels (**Figure 7**). Second, we found support for a non-strict hierarchical relationship between task and motor levels. Third, we provided some evidence that motor and sequence levels can have a non-strict hierarchical relationship. Finally, we did not find clear evidence that motor and abstract sequences formed an integrated construct with practice. Together, these findings provide insight about the mixed hierarchical relationships between levels in abstract task sequences under conditions that more closely resemble the complex and practiced sequential nature experienced in daily living.

**Figure 7.**
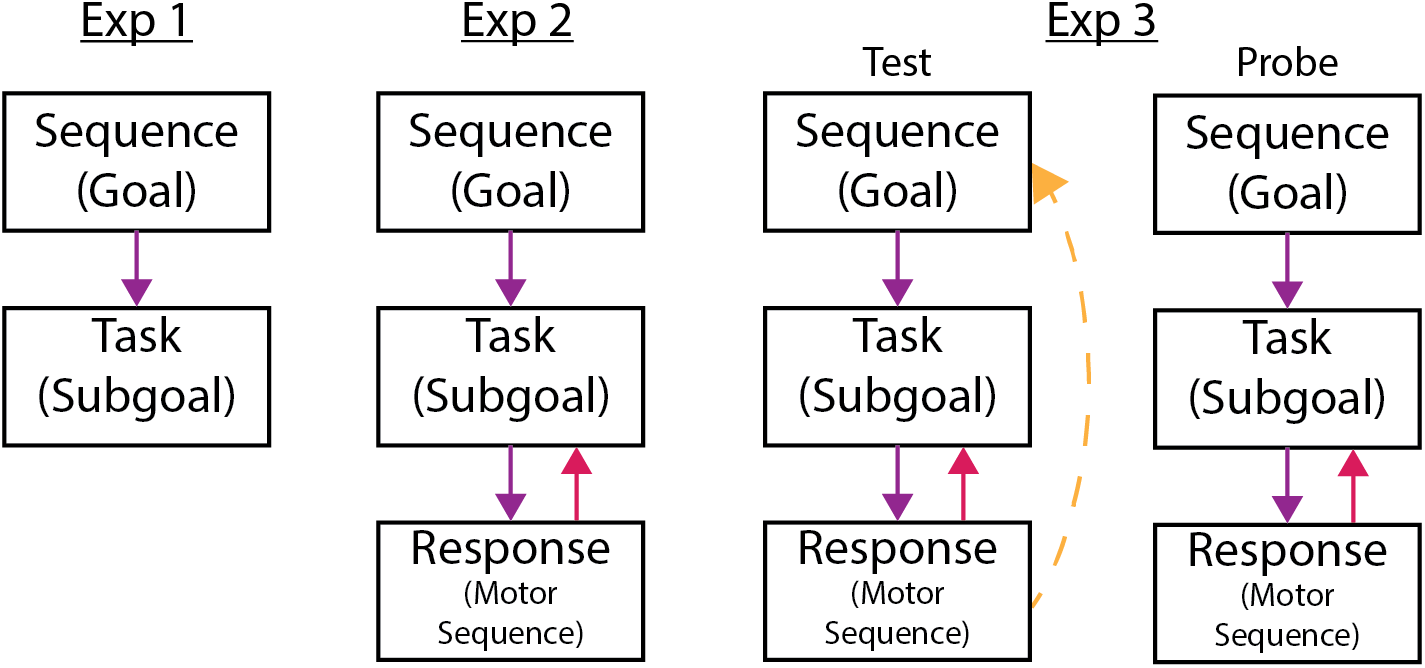
Summary of findings across experiments.

While practice effects on switch costs have been widely studied, the effects of practice on sequence initiation processes were unknown. Across all three experiments, practice at the sequence level specifically reduced initiation costs without affecting switch costs, indicating a strict hierarchical relationship between the sequence and task levels (**Figure 7**). Previous studies found a reduction, but not elimination, of switch costs with practice (Berryhill & Hughes, 2009; Stoet & Snyder, 2007; Strobach et al., 2012). Reductions in switch costs are hypothesized to reflect improvements in shifting attention to a new task set, inhibiting the irrelevant task set, or retrieving a new goal state (Hirsch et al., 2018; Sabah et al., 2019). Similarities between switch and initiation costs make it intuitive that initiation costs, like switch costs, may be reduced with practice. However, we did not observe a reduction in switch costs as the result of practice alone. The simplest explanation for the lack of effect on switch costs is that practice at the sequence level may have had uniform benefits at the task level such that there were no differences between the conditions at test. In other words, participants were able to generalize practice with task switching across sequence conditions. This possibility is supported by previous work that indicates that task switching practice effects generalize to cognitive control tasks that involve similar processes (Sabah et al., 2019) and the fact that sequence initiation costs are hypothesized to reflect task set reconfiguration that occurs at the beginning of each sequence (Schneider & Logan, 2006). This explanation raises the possibility that despite the strict hierarchical structure, similar processes could play a role in, and benefit from, practice at multiple levels of hierarchical representation. Additionally, it is possible that there was not sufficient practice to induce a change, or practice may need to occur specifically at the task control level to induce a change. While they cannot be ruled out, these options are less likely due to the amount of practice participants had, particularly in Experiments 2 and 3.

The nature of the hierarchical structure between the sequence and motor levels was less apparent. While Experiment 2 and the probe phase of Experiment 3 provided evidence that there was a strict relationship between the sequence and motor levels, the Experiment 3 test phase provided evidence that there was a non-strict relationship. Follow-up analyses comparing the test phases of Experiments 2 and 3; Experiment 2 test and Experiment 3 probe phases; and Experiment 3 test and probe phases were not consistent. These results are difficult to interpret and suggest that the nature of the relationship could be dependent on the specific context. A numerically greater number of participants in Experiment 3 relative to Experiment 2 noticed patterns in the motor responses for the sequence conditions. Further, the probe manipulation in Experiment 3 could have disrupted awareness of the motor sequences, causing results from the probe section to resemble those of Experiment 2. These observations suggest that awareness may influence hierarchical relationships, as awareness speeds reaction times in motor sequence execution (Wong et al., 2015). Though we addressed this question by comparing the putatively aware and unaware groups across the experiments, the results were ambiguous and did not point to a clear role of awareness. There are many possible explanations for this ambiguity, including that we did not directly manipulate awareness, and that the level of awareness may not have been sufficient to observe consistent effect. Further work is necessary to examine the role of awareness in how motor sequences affect abstract task sequence execution.

In contrast, there was consistent support for a non-strict hierarchical relationship between the motor and task levels (**Figure 7**). The reduction in switch costs in the Motor Familiar sequences were evident across both Experiment 2 and 3. Further, switch costs at the task and motor levels were not additive, but rather interacted such that a congruency effect was evident between switching and repeating across task and motor trial types. This finding replicates previous studies that examined interactions between task and response level information without the explicit inclusion of sequence-level information (Kikumoto & Mayr, 2020; Korb et al., 2017; Mayr & Bryck, 2005). The current results extend this finding and suggest that the interaction of the task and motor levels is a consistent feature of how tasks are performed, regardless of the overarching hierarchical structure.

Given that the addition of embedded motor sequences facilitated the execution of abstract task sequences and formed a non-strict representation between the motor and task levels, we designed the probe blocks in Experiment 3 to examine if and how these representations may be integrated. Previous task switching work has documented task-motor conjunctive representations suggesting that task levels can form integrated representations (Kikumoto & Mayr, 2020; Korb et al., 2017; Mayr & Bryck, 2005). Further, results from dual-task paradigms indicate that humans can integrate across simultaneously occurring sequential information and that this integration can facilitate learning (Cock & Meier, 2013; Schmidtke & Heuer, 1997; Weiermann et al., 2010; Weiermann & Meier, 2012). Together, this evidence suggests that humans are able to integrate information across task levels and that this type of integration might facilitate behaviors. We hypothesized that if there was an integrated representation, then disrupting the relationship between abstract task sequences with embedded motor sequences in the probe phase would cause task performance to be degraded, and that adding motor sequences to other abstract sequences would not be faciliatory. We did not find evidence for a degradation in performance, but instead found that incorporating embedded motor sequences into different abstract task sequences was faciliatory. Thus, we did not find evidence to support that there is an integration of representation between the abstract task and motor sequences, as they could be added and subtracted without disrupting the main patterns of results.

The offset between the abstract task sequences and the motor sequences may have discouraged an integrated representation. Support for this idea stems from the dual-task literature. The Motor Familiar condition in Experiments 2 and 3 could be conceptualized as a dual-task paradigm if the abstract sequence is considered as one “task” and the motor sequence as another “task.” As in our experiments, participants in dual-task paradigms were unable to reproduce the embedded motor sequences (Heuer et al., 2001; Schumacher & Schwarb, 2009; Schwarb & Schumacher, 2012). Participants in dual-task paradigms benefited when both tasks (i.e., abstract and motor) contained sequential information (Cock & Meier, 2013; Heuer et al., 2001), but less so, or not at all, when the two tasks contained repetitive sequences of different lengths (Cock & Meier, 2013; Schmidtke & Heuer, 1997; Weiermann & Meier, 2012). Because participants in the current experiments realized a benefit in performance with sequences of different lengths, these results suggest that the sequence offset itself may not be responsible for a lack of integration. Other methodological differences such as the length of the sequences or the explicit instruction of the abstract task sequence could also explain these differences. Therefore, the relationship between sequence lengths at different levels of the hierarchy and the potential for integration should be explored in future experiments. Our experiments have introduced a novel task paradigm that can be used as a tool for such investigation.

There are two limitations of the current design that are mitigated by the probe block manipulation in Experiment 3. First, we did not include a condition in the test blocks where novel, unpracticed abstract task sequences were performed with embedded motor sequences. This design choice was due to our focus on the differential effects of the embedded motor sequences on the abstract task sequences and to maintain a reasonable number of trials that participants could perform in a single session and maintain a within-subjects design. Therefore, interactions between the motor and superordinate levels could have been observed because they were practiced together. The Familiar Plus condition mitigates this concern because the abstract task sequence was not practiced with an embedded motor sequence, yet, participants benefited from the addition of the motor sequence. This result suggests that interaction between the hierarchical levels is not a direct result of the sequences being practiced together.

Second, the facilitation of Motor Familiar sequences may have resulted from participants memorizing specific task and response associations. Though there were design features that discouraged that situation, such as the offset between the abstract task and motor sequences and the 20 trials between each time the sequences aligned, this possibility remained. Conditions in the probe phase of Experiment 3 mitigated this possibility. We observed a facilitation in reaction times in the Familiar Plus condition despite that participants had no opportunity to associate the specific abstract tasks with the embedded sequential motor responses. This result provides further evidence that the performance benefits of adding a motor sequence were not due to participants memorizing a longer stimulus sequence.

While we did not specifically examine the mechanisms by which embedded motor sequences facilitated abstract task sequence performance, the results are consistent with a number of possibilities. First, conceptualized as a dual-task, the experimental paradigm could have led to a parallel race between responses at the task and motor levels (e.g., Rowe et al., 2010). When task selection was delayed (i.e., switch trials), implicit knowledge of the motor sequence could facilitate a faster reaction time compared to trials with no motor sequence. Similarly, the presence of iterating sequences at the sequence and motor levels allows participants to know, explicitly or implicitly, about upcoming information. As discussed before, the combination of a task, specified by the task sequence, and a response, specified by the motor sequence, dictates the relevant stimulus parameter for an upcoming trial. Thus, it is possible that these convergent streams of information allow participants to make better predictions about upcoming stimuli and thus facilitate choices.

Another possibility is derived from automatic control theory (Logan, 2018). In the execution of practiced motor sequences, this theory posits that the effect of practice is to offload the execution of the motor actions from the working memory system to the motor system. Therefore, control costs are reduced due to a reduction in the use of a common resource, as opposed to the specific control processes themselves. While Logan’s (2018) theory provides an account of the execution and control of very well-learned motor skills (e.g., typing), it leaves open the question of skills that have a more intermediate level of practice, as well as the process of acquisition. We provide evidence of selective improvement of control costs at the task and abstract task sequence levels with practice and embedded motor sequences. This theory is further supported by recent work showing that other hierarchical control structures show improvements across levels with practice (Yokoi & Diedrichsen, 2019). Therefore, with the extent of practice that exists in daily living, it is possible that such control processes could be further optimized to become automatic and potentially rely even less on working memory resources. Explicit tests with extended practice and of automaticity and working memory would be necessary to further this theory beyond the initial evidence we provide here.

An important avenue of future research will be to disentangle the potential control mechanisms necessary for both abstract task and motor sequences when they are extensively practiced, as they commonly are in daily life. How the brain supports their execution could provide important insight regarding which specific processes are facilitated, and whether abstract task sequences and motor sequences use common resources. The rostrolateral prefrontal cortex (RLPFC) is necessary for the execution of abstract task sequences and is among a network of areas that shows dynamics that may be unique to sequential control (Desrochers et al., 2015, 2019). Motor sequence acquisition and performance is supported by a network of areas that include subcortical areas, such as the striatum and the cerebellum, and motor cortical areas (Keele et al., 2003; Robertson, 2007; Wiestler et al., 2014) as well as the medial temporal lobe and prefrontal cortex (Destrebecqz et al., 2005; Schendan et al., 2003). This network may overlap with those observed in abstract task sequences. Increasing our understanding of the overlap of these systems will necessitate examining the simultaneous performance of abstract and motor sequences, and we have presented a novel paradigm that is capable of addressing these and similar questions.

In conclusion, these studies present new evidence that practice and embedded motor sequences facilitate abstract task sequence execution. We provide new insight into the interrelations between hierarchical levels (goal, sub-goal, and motor) common in many task paradigms by replicating and extending these results to the context of more abstract task sequences. These findings suggest that the relationship between levels may be specific to the context and highlight the necessity of studying complex hierarchical structures together, rather than in isolation. Overall, these studies demonstrate that the facilitation of control costs at the goal and sub-goal levels are possible mechanisms for efficient execution of complex tasks in daily life.

## Acknowledgements

The authors would like to thank David Badre and Kathryn Graves for work and discussions on an earlier, related project, and the development of this project. We would like to thank Matthew Maestri, Sarah Master, Victoria Flagg, Eojin Choi, Jeff Mercurio, Guillaume Pagnier, and Gabriela Batista for their assistance with data collection. We also thank members of the Badre and Desrochers Labs for many helpful discussions during the preparation of this manuscript. Research reported in this publication was supported by the Brown University UTRA program (J.E.T.), NIDA F32 DA045451 (T.H.M.), and Brown University Department of Neuroscience Connors Fellowship (T.H.M.), NIGMS COBRE (P20GM103645, T.M.D.), NSF EPSCOR Track-2 (1632738, T.M.D.), and the Robert J. and Nancy D. Carney Institute for Brain Science (Innovation Award, T.M.D.).

## Data Availability

The raw data supporting the conclusions of this manuscript may be made available by the authors, without undue reservation, to any qualified researcher. These studies were not preregistered.

## Notes

### Competing Interest Statement

The authors have declared no competing interest.

### Summary of Updates

Reframed results in terms of hierarchy. Added summary figures. Added analyses to test hierarchy hypotheses.

